# Mutational spectra analysis reveals bacterial niche and transmission routes

**DOI:** 10.1101/2022.07.13.499881

**Authors:** Christopher Ruis, Aaron Weimann, Gerry Tonkin-Hill, Arun Prasad Pandurangan, Marta Matuszewska, Gemma G. R. Murray, Roger C. Lévesque, Tom L. Blundell, R. Andres Floto, Julian Parkhill

## Abstract

As observed in cancers, individual mutagens and defects in DNA repair create distinctive mutational signatures that combine to form context-specific spectra within cells. We reasoned that similar processes must occur in bacterial lineages, potentially allowing decomposition analysis to identify disrupted DNA repair processes and niche-specific mutagen exposure. Here we reconstructed mutational spectra for 84 clades from 31 diverse bacterial species, assigned signatures to specific DNA repair pathways using hypermutator lineages, and, by comparing mutational spectra of clades from different environmental and biological locations, extracted reproducible niche-associated mutational signatures. We show that mutational spectra can predict general and specific bacterial niches and therefore reveal the site of infection and types of transmission routes for established and emergent human bacterial pathogens.

**One sentence summary:** Variable mutagen exposure and DNA repair drive differential mutational spectra between bacteria and enable niche inference

---

Work using human cells and tissues has demonstrated that mutagens induce highly specific context-dependent patterns of base substitutions termed mutational signatures, which combine to form a mutational spectrum (*1-6*). However, these patterns are compounded with the signatures of endogenous mutations and DNA repair, which also exhibit specific mutational signatures (*7-9*).

Reconstructing the set of mutations and signatures within cancers has enabled inference of the drivers of tumourigenesis (*1, 2, 7*). We therefore reasoned that reconstructing mutational spectra in bacteria, differentiating them into different signatures, and correlating these with known DNA repair defects and environmental exposures, should allow the association of specific DNA signatures with bacterial niches. These signatures could then be used to predict niche or infection sites and to identify defects in DNA repair when niche is known. To test this, we undertook the first large-scale comparison of mutational spectra and their underlying signatures across bacteria, correlating the results with DNA repair pathways and niche.

We used whole genome sequence alignments and phylogenetic trees to reconstruct single base substitution (SBS) mutational spectra of 84 phylogenetic clades from 31 diverse bacterial species, implemented in a specifically-developed open-source bioinformatic tool, MutTui (**fig. S1; fig. S2; table S1; table S2**; **Supplementary Methods**). SBS spectra were rescaled by genomic nucleotide composition to enable direct comparison between bacteria. We find that such spectra are highly diverse, both in the nucleotide mutations themselves and their surrounding context (**Fig. 1; fig. S2**). However, several generalisable properties could be identified. We found that transition mutations are more common than transversion mutations (*10*) in all cases (ranging from 52-55% in *Klebsiella pneumoniae* to >90% in *Campylobacter jejuni*; **fig. S2**). Cytosine to thymine (C>T) was typically the most common mutation type identified (in 69 of 84 SBS spectra examined), potentially due to cytosine deamination (*11*). Genomic G+C content exhibits a negative correlation with proportion of C>A/T mutations but a positive correlation with proportion of C>G mutations (**fig. S3**). Finally, transition mutations exhibit enriched context specificity compared to transversion mutations while several contextual mutations are significantly elevated across datasets (**fig. S4**). UMAP clustering revealed groups of similar SBS spectra across bacterial clades (**Fig. 1**). We observe a strong correlation between phylogenetic relatedness and spectrum similarity (Tukey HSD corrected ANOVA *P* < 0.001; **fig. S5**) and spectra are typically conserved across highly-related clades where there has likely been no change of niche or DNA repair capacity (**fig. S6**).

**Fig. 1.**
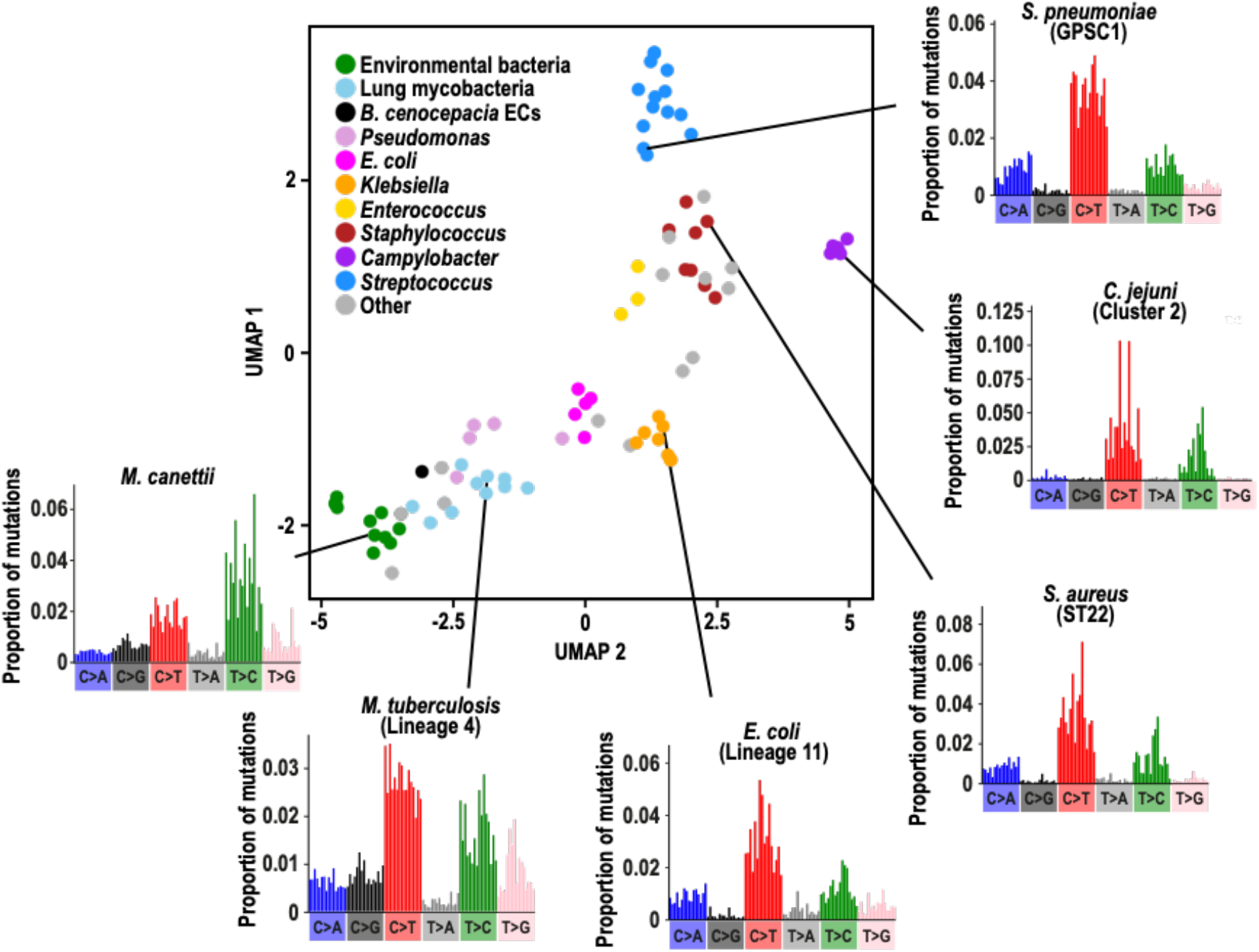
Clustering of bacterial SBS spectra. UMAP clustering based on contextual mutation proportions within the 84 SBS spectra across 31 bacterial species. Selected groups are coloured. The environmental bacteria label includes *Burkholderia pseudomallei* and known environmental *Mycobacteria*. Example SBS spectra are shown for selected groups.

We reasoned that bacterial SBS spectra can be decomposed into combinations of mutational signatures, each driven by distinct defects in DNA repair or by endogenous processes or specific mutagens, as has previously been achieved for cancer-associated mutations (*1, 3, 7*). Therefore, we first extracted mutational signatures associated with distinct DNA repair pathways by calculating SBS spectra of 50 naturally-occurring hypermutator lineages across four bacterial species (**Fig. 2A**). By identifying the genes most likely responsible for hypermutation, we were able to attribute mutational signatures to defects in 11 DNA repair genes that function in mismatch repair (MMR), base excision repair (BER), or homologous recombination (HR) (**Fig. 2B-D; Supplementary Methods**).

**Fig. 2.**
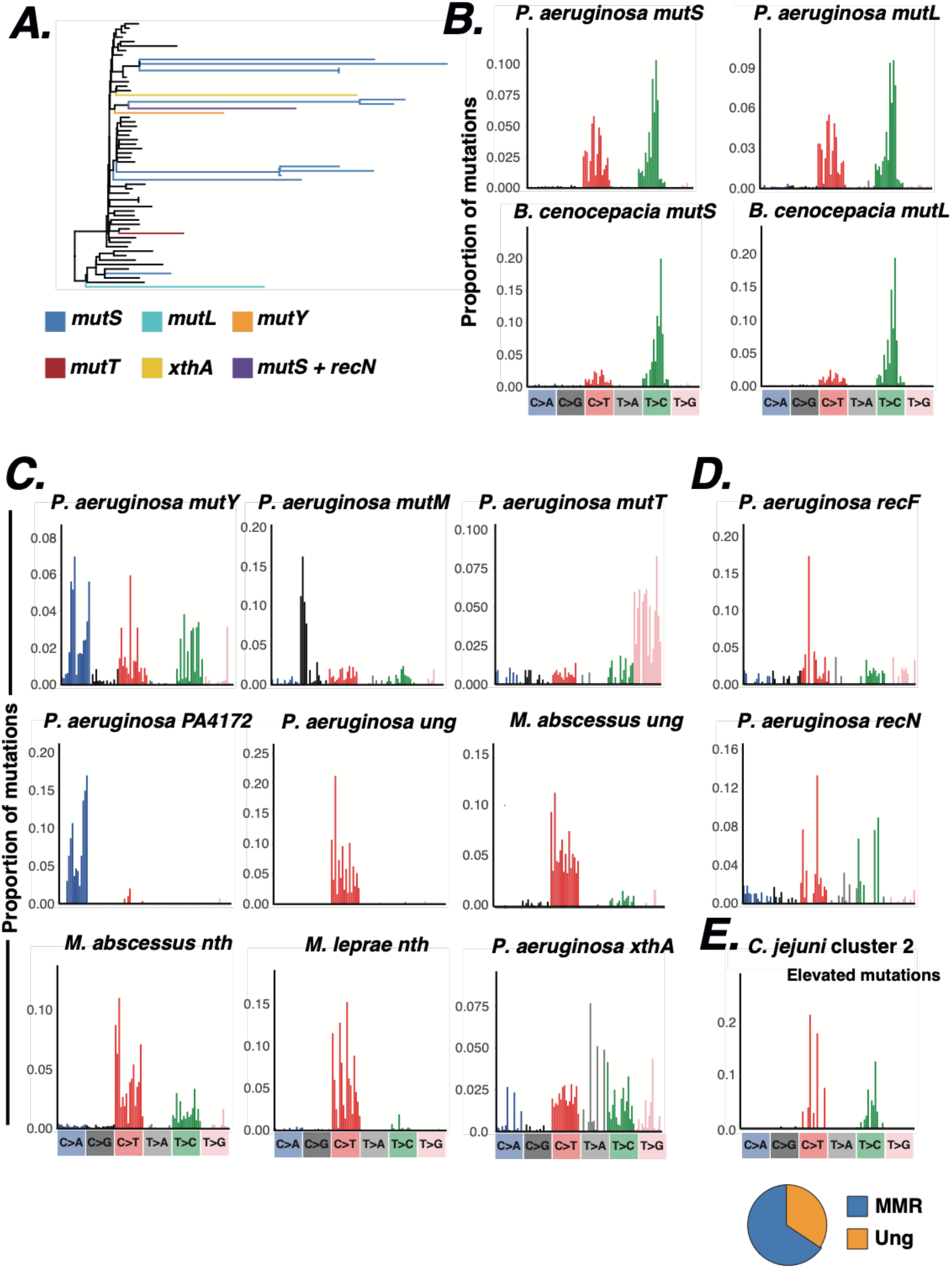
Mutational signatures associated with DNA repair genes. (**A**) Example *P. aeruginosa* phylogenetic tree (ST274) showing hypermutator branches and the inferred responsible genes. Hypermutator branches were identified based on branch length and the ratio of transition and transversion mutations. Responsible genes were identified as DNA repair genes exhibiting a mutation on the long phylogenetic branch or ancestral branch. Black branches are background non-hypermutator branches that did not contribute to hypermutator spectra. (**B**) Mutational signatures associated with MMR genes. (**C**) Mutational signatures associated with BER genes. (**D**) Mutational signatures associated with genes involved in homologous recombination. (**E**) Top panel shows the mutations elevated in *C. jejuni* cluster 2 compared with *E. coli* lineage 34, calculated by subtracting each respective mutation proportion in the SBS spectra. The pie chart shows the proportion of mutations elevated in *C. jejuni* cluster 2 that are assigned to each bacterial DNA repair gene signature in a decomposition analysis.

Mutations of MMR genes result in high levels of context-specific C>T and T>C mutations (**Fig. 2B, fig. S7**) (*1, 8, 12*) which likely represent the error profile of DNA Polymerase III that is usually repaired by functional MMR. While context specificity is highly similar between species, the relative rates of C>T and T>C differ between *Pseudomonas aeruginosa* and *Burkholderia cenocepacia* (**Fig. 2B; fig. S8**), likely reflecting distinct polymerase error profiles (a possibility supported by structural modelling analysis; **fig. S9**).

Mutations in distinct base excision repair (BER) components results in characteristic gene-specific patterns (**Fig. 2C**), as expected from the diverse repair functions of proteins within this pathway (*11*). We identified *P. aeruginosa* hypermutators for each component of the GO repair pathway (*mutT, mutY* and *mutM*) that prevents 8-oxoguanine (8-oxo-G)-induced mutations (*13*). Mutation of *mutT*, whose product degrades 8-oxo-G monomers to prevent their incorporation into DNA (*13*), results in non-specific T>G mutations (**Fig. 2C**), suggesting incorporation of 8-oxo-G opposite adenine is context-independent. Conversely, mutation of *mutY* which excises adenine opposite 8-oxo-G (*13*), results in C>A mutations predominantly in CpCpN and TpCpN contexts (**Fig. 2C**), indicating context-specific mutation of incorporated guanine to 8-oxo-G. This likely represents the pattern of reactive oxygen species (ROS) damage, of which 8-oxo-G is a major mutagenic lesion (*9*). The C>A contexts differ between the *P. aeruginosa mutY* signature and human cell signatures of ROS exposure (*5*) and knockout of either the *mutY* homologue or OGG1 (*7, 9*) (**fig. S10**), suggesting differential repair of these lesions by other proteins. Mutation of *mutM* results in C>G mutations in ApCpN contexts (**Fig. 2C**). While the mechanism of C>G mutations is unclear, the lack of C>A mutations in *mutM* knockouts is potentially due to functional MutY being sufficient to repair mutagenic 8-oxo-G lesions (*14*). We additionally identify PA4172 in *P. aeruginosa* whose knockout exhibits C>A mutations in CpCpN and TpCpN contexts similar to *mutY* (Pearson’s *r P* < 0.001; **Fig. 2C; fig. S10**), suggesting that its product may similarly repair mutagenic 8-oxo-G lesions.

Disruption of *ung*, whose product removes uracil from DNA (*11*), results in similar patterns of context-specific C>T mutations in *P. aeruginosa* and *Mycobacterium abscessus* (Pearson’s *r P* < 0.001; **Fig. 2C; fig. S11**). This bacterial signature exhibits subtle contextual differences compared with *ung* knockout in human cells (*9*), particularly through enriched mutations in NpCpG contexts (**fig. S11**), suggesting differential patterns of uracil incorporation in humans and bacteria.

Mutation of *nth*, whose product Endonuclease III removes damaged pyrimidines, results in C>T mutations in multiple *Mycobacteria* species and human cells (*8*) but with different context specificity (Pearson’s *r P* > 0.05; **Fig. 2C; fig. S12**). Disruption of the apurinic-apyrimidinic (AP) endonuclease *xthA* results in mutations in multiple specific contexts (**Fig. 2C**), particularly transversions in [C,G,T]p[C,T]pG contexts, indicating repair of a broad range of specific lesions. Finally, hypermutators resulting from mutation of the homologous recombination pathway components *recF* and *recN* exhibit context-specific transition mutations (**Fig. 2D**). Recombination is known to drive GC-biased gene conversion (*15*) and this may contribute to this signature.

We subsequently tested whether we could detect differences in SBS spectra from bacteria with different repair capabilities occupying a similar niche (and therefore exposed to similar sets of niche-specific mutagens). Decomposition analysis showed that almost all mutations elevated in *C. jejuni* compared with the gastrointestinal *Escherichia coli* lineage 34 (*16*) can be explained by a failure to repair deaminated cytosines and a lack of MMR (**Fig. 2E; fig. S13)**; pathways which are known to be absent in *C. jejuni* (*17*). Our results indicate that differences in DNA repair can be inferred by comparing bacteria from a similar niche.

We then proceeded to extract further bacterial signatures through a decomposition analysis employing nonnegative matrix factorisation (NMF) (*18, 19*) on SBS spectra datasets from a range of species and genera (**table S3**). We extracted 33 SBS signatures and collapsed these into a final set of 24 (named with the prefix Bacteria_SBS) by combining highly similar signatures (with cosine similarity of 0.95 or greater) (**fig. S14; table S4**). The extracted signatures exhibit divergent base mutations and contexts and are differentially present across bacteria (**fig. S14**), supporting differential activity of mutagens and repair between clades. An exception to this pattern was signature Bacteria_SBS15 that was extracted from the *Staphylococcus* genus dataset and the *Enterococcus faecalis, Streptococcus pneumoniae* and *Streptococcus agalactiae* species datasets (**fig. S14**), indicating broad distribution across Bacillota. As these bacteria inhabit different niches, this signature likely represents phylum-specific endogenous mutations and/or DNA repair profiles.

We next explored the influence of pathogen niche on mutational spectrum, focussing on *Mycobacteria* and *Burkholderia*, genera that contain both clades that are transmitted from person-to-person and clades that are acquired from environmental sources (*20, 21*). We find that known lung and environmental clades cluster separately based on SBS spectrum composition (**Fig. 3A**). Spectrum subtractions consistently revealed elevated C>A and C>T mutations in lung bacteria and higher levels of T>C in environmental bacteria (**Fig. 3B, C**). Lung and environmental bacteria additionally exhibit different contextual patterns within C>A and T>C mutations (**fig. S15**). Decomposition of niche-specific mutations from subtracted spectra using known human mutagen signatures suggests that higher C>A in lung bacteria is likely driven by tobacco smoke (*2*) (found in human but not animal infecting lung clades) and exposure to reactive oxygen species (ROS), while higher T>C within the environment is probably caused by exposure to alkylating agents and nitro-polycyclic aromatic hydrocarbons (*5*) (**Fig. 3D**), mutagens known to be present in the environment (*5*). It is also possible that the long-term evolutionary selection towards GC richness seen in some bacterial genomes (*22*) may contribute to the observed environmental signature.

**Fig. 3.**
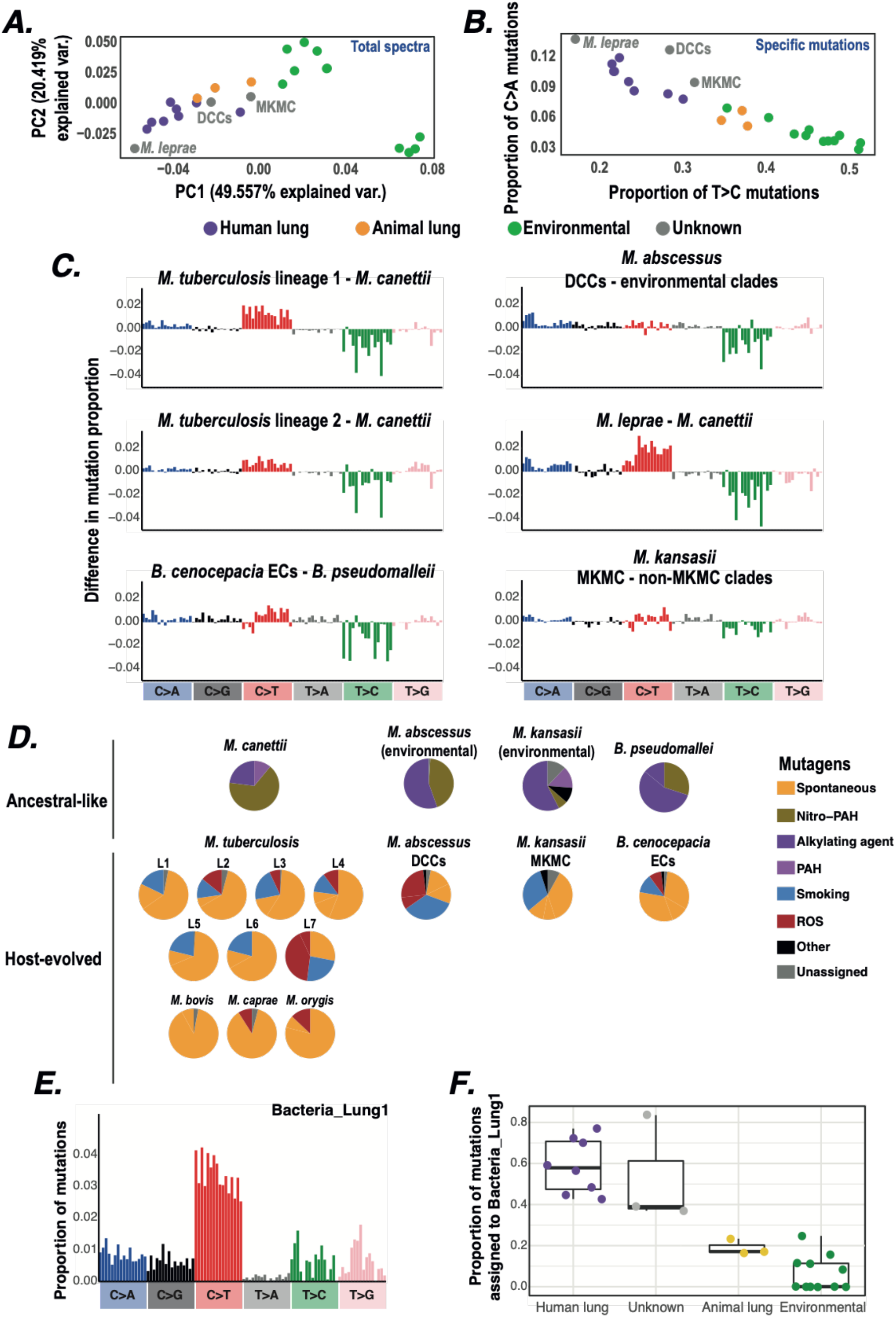
Comparison of mutational spectra between lung and environmental niches. (**A**) Principal component analysis on mutation proportions in the SBS spectra across *Mycobacteria* and *Burkholderia*. Axes labels include the inferred proportion of variance each principal component describes. Points are coloured by niche; clades with a previously unknown niche are labelled. Environmental includes *B. pseudomallei* and known environmental clades of *Mycobacteria*. (**B**) Comparison of the proportion of T>C and proportion of C>A mutations in *Mycobacteria* and *Burkholderia* SBS spectra, coloured as in **A**. (**C**) Subtraction of mutation proportions in SBS spectra between closely related bacterial clades. Each comparison subtracts the SBS spectrum of a known environmental clade from the SBS spectrum of a clade either known to reside within the lung or with an unknown niche. (**D**) Decomposition of mutational spectra into their underlying components. Only mutations elevated within the respective clade compared to a closely related clade in a different niche were included. Known environmental clades were decomposed into the set of previously extracted environmental mutagen signatures (*5*) while known lung clades and clades with unknown niche were decomposed into the set of previously extracted lung signatures from human data. *B. cenocepacia* ECs: *B. cenocepacia* epidemic clones. Nitro-PAH: nitro-polycyclic aromatic hydrocarbons; PAH: polycyclic aromatic hydrocarbons; ROS: reactive oxygen species. (**E**) Composition of signature Bacteria_Lung1 extracted from NMF decomposition of *Mycobacteria* and *Burkholderia* SBS spectra. (**F**) The proportion of mutations within each *Mycobacteria* and *Burkholderia* SBS spectrum assigned to signature Bacteria_Lung1.

We further examined niche signatures through a targeted NMF decomposition of the *Mycobacteria* and *Burkholderia* spectra and were able to extract a lung-associated mutational signature consisting of multiple mutation types that we term Bacteria_Lung1 (**Fig. 3E, F**). Building on these different patterns, we developed a set of leave-one-out classifiers and found that they were able to robustly predict known lung or environmental niche based on either SBS spectrum, proportion of the six mutation types or cosine similarity between SBS spectra (**table S5**; **fig. S16**).

Due to their success in predicting known niches, we next used SBS spectra to infer niche for several *Mycobacteria* clades where this was previously unknown. We find strong evidence that *Mycobacterium leprae* and the dominant circulating clones (DCCs) of *M. abscessus* (*23*) replicate within the lung as they: cluster with known human lung bacteria based on their SBS spectrum (**Fig. 3A**); exhibit lung-like contextual patterns of C>A and T>C mutations (**fig. S15**); exhibit high levels of C>A and low levels of T>C (**Fig. 3B, C**); and exhibit signature Bacteria_Lung1 at similar levels to known lung bacteria (**Fig. 3F**). These observations suggest human-to-human lung transmission and are supported by reports that *M. leprae* can replicate within human alveolar epithelial cells *in vitro* and can infect mouse lung macrophages and epithelial cells during *in vivo* challenge (*24*), and that the *M. abscessus* DCCs have spread through global transmission chains involving Cystic Fibrosis (CF) and non-CF individuals (*20, 23*).

The *Mycobacterium kansasii* main cluster (MKMC) causes the majority of *M. kansasii* infections (*25*) and exhibits characteristics of both lung and environmental spectra. Specifically, the MKMC exhibits lung-like C>A patterns but environmental-like T>C patterns (**fig. S15**) and is therefore intermediate between known human lung and environmental spectra in the SBS clustering (**Fig. 3A**) and C>A vs T>C comparison (**Fig. 3B**). Together, these results suggest that the MKMC is exposed to both lung and environmental mutagens and therefore replicates within (and is potentially acquired from) both niches.

We next examined multiple niches within the same host by comparing human *Salmonella* lineages that cause enteric infection with those that have adapted to cause invasive disease (**table S1 &, S6**). The SBS spectra cluster by niche (**Fig. 4A**; association index *P* < 0.001), rather than by phylogeny. While we could not identify clear and conserved differences between SBS spectra in the different niches by eye (**fig. S17**), we were again able to develop classifiers that could robustly predict enteric or invasive niche (**table S5; fig. S16**), suggesting that even subtle spectrum differences can be sufficient to reliably distinguish niche.

**Fig. 4.**
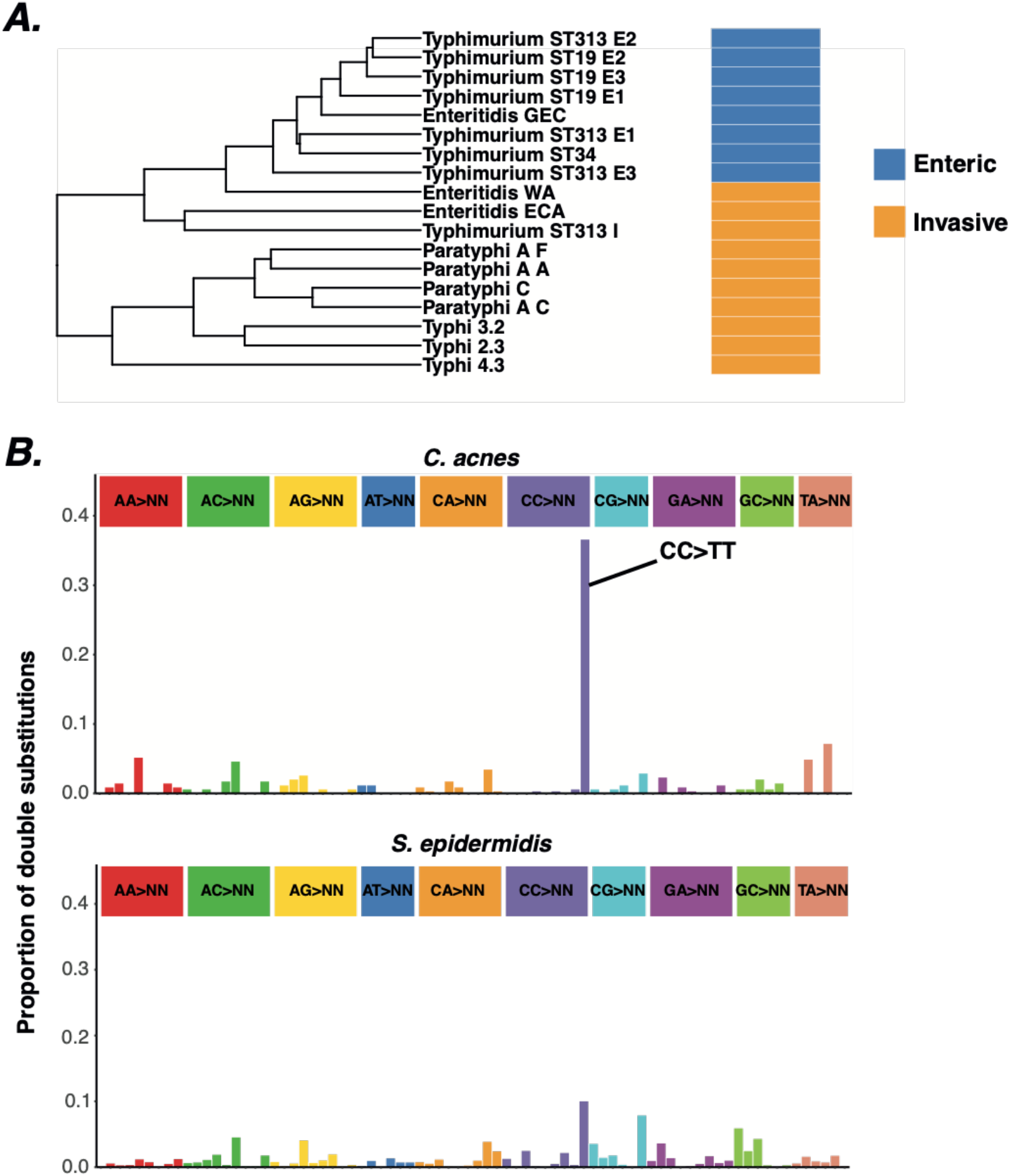
Comparison of mutational spectra between niches. (**A**) Hierarchical clustering of mutation proportions in salmonella SBS spectra labelled by niche. The enteric clades within Typhimurium ST19 and ST313 were split into three subclades for this analysis, labelled E1-E3. Typhimurium ST313 also contains an invasive clade labelled I. GEC - Global Enteric clade; WA - West Africa clade; ECA - East and Central Africa clade. Three genotypes of Paratyphi A were included - A, C and F. (**B**) Comparison of double substitution spectra between *C. acnes* and *S. epidermidis* (phylogenetic groups A-C combined). CC>TT is indicated in the *C. acnes* spectrum and is a classic signature of exposure to UV light (*5*).

Finally, we tested whether mutational spectra could distinguish sub-niches within the same host niche. We find a high level of CC>TT double mutations characteristic of UV-light damage (*5*) in the pan-skin bacterium *Cutibacterium acnes* that is not present in *Staphylococcus epidermidis* which preferentially inhabits moist (*26*), and therefore less sun-exposed, skin sites (**Fig. 4B**). Together, these results demonstrate that mutational spectra can predict bacterial niche with very high levels of spatial resolution.

In conclusion, we show that we can reconstruct mutational spectra from bacterial phylogenies and decompose these into specific signatures. We can ascribe some of these signatures to defects in DNA repair pathways, and others to exposure to location-dependent mutagens which can be used to predict the niche in which bacteria replicate and infer transmission routes. We anticipate that identification of signatures at different levels in bacterial phylogenies will identify ancestral niches and therefore sources of emergent human pathogens, reveal routes of acquisition of infection permitting targeted interventions, and provide a mechanism to monitor pathogenic evolution and host adaptation. We envisage that mutational spectra analysis could be applied to viruses and parasites, enabling similar predictions.

## Acknowledgements

The Authors would like to thank all researchers who helped to obtain published datasets used in this study, including Uzma Basit Khan, Christopher Beaudoin, Sophie Belman, Stephen Bentley, Sebastian Bruchmann, Jessica Calland, Claire Chewapreecha, Jukka Corander, Dorota Jamrozy, Anna Kaarina Pöntinen, Noémie Lefrancq, Stephanie Lo, Neil MacAlasdair, Samuel Sheppard, Andries van Tonder and Lucy Weinert.

## Author contributions

Conceptualization: CR, RAF, JP

Methodology: CR, GTH

Investigation: CR, AW, APP, MM, GGRM, RCL

Visualization: CR, APP

Funding acquisition: TB, RAF, JP

Project administration: RAF, JP

Supervision: TB, RAF, JP

Writing - original draft: CR, RAF, JP

Writing - review & editing: CR, AW, GTH, APP, MM, GGRM, RCL, TB, RAF, JP

## Data and materials availability

All data and code used for data analysis is available at https://github.com/chrisruis/Mutational_spectra_data. The MutTui pipeline used to reconstruct pathogen mutational spectra is available at https://github.com/chrisruis/MutTui.

## Supplementary Materials

Materials and Methods

Figs. S1 to S17

Tables S1 to S8

References (27-36)

