## Supplementary Material and Figures for "Mutational spectra analysis reveals bacterial niche and transmission routes"

**This PDF file includes:**

Materials and Methods
Figs. S1 to S17

**Other Supplementary Materials for this manuscript include the following:**

Tables S1 to S8

### Materials and methods

Dataset sources and reconstruction of phylogenetic trees

We collated published whole genome sequencing datasets from 84 phylogenetic clades across 31 bacterial species (**table S1,** accession numbers listed in **table S7**). Datasets were obtained either from public databases as FASTQ files or genome assemblies, or from study Authors as whole genome sequence alignments or post-recombination removal variable sites alignments (**table S1**). Where datasets were obtained as genome assemblies, they were initially shredded to FASTQ files containing 100 base pair reads with a 350 base insert size and depth 40 using Fastaq v3.17.0 (https://github.com/sanger-pathogens/Fastaq). Sequencing reads from obtained FASTQ files and shredded assemblies were mapped against a clade-specific reference genome **(table S1**) using the multiple_mappings_to_bam pipeline v1.6 (https://github.com/sanger-pathogens/bact-gen-scripts) with BWA-MEM as the aligner. Recombination was removed from mapped alignments and whole genome sequence alignments obtained from study Authors using Gubbins v2.4.1 (*27*). Maximum likelihood phylogenetic trees were reconstructed from post-recombination removal variable sites alignments for all datasets using RAxML v8.2.12 (*28*) with the general time reversible (GTR) model of nucleotide substitution and gamma rate heterogeneity with four gamma classes.

Reconstruction of bacterial mutational spectra

We reconstructed mutational spectra using MutTui v1.1.10 (https://github.com/chrisruis/MutTui) which employs the variable sites alignment, phylogenetic tree and reference genome for each dataset. Mutations are reconstructed onto the phylogenetic tree using treetime v0.8.1 (*29*) which enables identification of the direction of each mutation. It was not possible to identify suitable outgroups to root phylogenetic trees for many datasets and we therefore used midpoint rooted phylogenetic trees. The surrounding nucleotide context (defined as the nucleotide immediately 5' and nucleotide immediately 3') of each mutation is inferred from the genome sequence at the start of the respective phylogenetic branch, identified by incorporating substitutions between the root of the tree and the start of the branch into the genome sequence at the root of the tree. We therefore infer the context of each mutation at the time it occurred. The mutational spectrum is constructed by counting the numbers of each contextual mutation across the clade. Single nucleotide mutations are included in the single base substitution (SBS) spectrum, mutations at two adjacent genome positions on the same phylogenetic branch are included in the double base substitution (DBS) spectrum and tracts of mutations at three or more adjacent genome positions are excluded. To account for differences in G+C content and triplet availability between clades, we rescale each spectrum by dividing the count of each contextual mutation by the frequency of the starting triplet in the clade reference genome.

For several datasets (**table S1**), the two phylogenetic branches that diverge immediately from the root of the tree represented a large proportion of the mutations in the dataset. We excluded mutations on these root branches in these cases as, due to the necessity to use midpoint rooted trees, their direction may not be inferred accurately and they would account for a large proportion of mutations in the spectrum. Several datasets exhibited evidence of hypermutator branches (**table S1**) which were excluded from the main clade SBS spectrum but split into separate SBS spectra based on the mutated gene (described in more detail below).

We used the *M. abscessus* SBS spectra we calculated previously where phylogenetic branches were divided into DCCs and non-DCCs (*23*). The *M. kansasii* phylogenetic tree contains both the MKMC clade and non-MKMC branches (*25*). We split these into separate SBS spectra by labelling branches in the phylogenetic tree as MKMC or non-MKMC which enables MutTui to extract a separate SBS spectrum for each group. The *Burkholderia pseudomallei* genome contains two chromosomes (*21*). We calculated the SBS spectrum of each chromosome separately to enable removal of recombination. Due to very high similarity between SBS spectra from chromosomes one and two in each group (cosine similarity >0.99 in each case), we used the chromosome one SBS spectra in further analyses. The *B. cenocepacia* SBS spectrum includes three epidemic clones whose SBS spectra were calculated separately and combined for further analyses.

Overall SBS spectra were compared using UMAP (*30*) based on the proportion of each of the 96 contextual mutations in each SBS spectrum. To examine the relationship between phylogenetic relatedness and overall spectrum similarity, we calculated the cosine similarity between all pairs of SBS spectra and split the comparisons into within-species, within-genus but different species, within-phylum but different genus and different phylum. The distributions of cosine similarities were compared between groups using two-way ANOVA with Tukey Honestly Significant Difference (HSD) correction.

We compared the degree of context-specificity between mutation types by calculating the variance of the contextual mutation proportions within each of the six mutation types in each of the 84 SBS spectra. The variance distribution between mutation types was compared using two-way ANOVA with Tukey HSD correction. Individual contexts within a mutation type were inferred to be significantly elevated or reduced if their median proportion within the respective mutation type across the 84 SBS spectra was more than 2.5 times the median absolute deviation outside the median of all context proportions in the mutation type.

#### Identification of DNA repair gene mutational signatures

We identified potential hypermutator lineages as very long branch lengths (either terminal branches or internal branches where each downstream branch is long) within phylogenetic trees across the 84 datasets; such lineages were identified in *P. aeruginosa*, *B. cenocepacia* and *M. leprae*. We additionally examined a broader *P. aeruginosa* dataset consisting of 18 sequence types and identified hypermutator lineages in this dataset. For each clonal cluster in this dataset, we compared the ratio of transition mutations to transversion mutations on each branch to the background distribution to identify candidate hypermutator branches (Fisher exact test, padj < 0.1). We only included branches in the background distribution that had at most 50 substitutions.The gene likely responsible for the hypermutation in each lineage was inferred through identifying DNA repair genes that exhibit a frameshift, insertion/deletion or nonsynonymous mutation on either the hypermutator branch or an upstream branch where each of the descendent branches are hypermutators. We identified the effects of these mutations using MutTui v1.1.10 applied independently to the branches containing the mutations.

Where a DNA repair gene was mutated on multiple branches, we calculated the SBS spectrum of the mutant as the mean mutational spectrum across branches. We excluded several *P. aeruginosa* branches that had both *mutS* and *mutL* mutations. In several cases, *mutS* and another DNA repair gene were mutated on the same branch; we here calculated the SBS spectrum of the other DNA repair gene by subtracting the mean *mutS* SBS spectrum from the branch SBS spectrum.

We additionally included two previously identified *M. abscessus* hypermutator lineages that arose within individual chronic pulmonary infections (*31*). The genes responsible for the hypermutation and the full set of mutations within the hypermutator lineages were previously inferred (*31*). We used MutTui v1.1.10 to identify the surrounding nucleotide context of each mutation from a closely related reference genome (*31*).

To compare the extracted bacterial DNA repair gene signatures with those previously calculated in human cells, we obtained COSMIC SBS signatures from https://cancer.sanger.ac.uk/signatures/sbs/ (date last accessed 24/06/2022) and gene knockout signatures from https://signal.mutationalsignatures.com/ (date last accessed 24/06/2022). We compared mutational patterns through a regression of the proportions of the 16 contextual mutations within the mutation type that is dominant within the respective gene signatures and applied a Benjamini-Hochberg correction on p-values from all comparisons.

Identification of defective DNA repair signatures in C. jejuni

To identify mutations that are likely the result of defective DNA repair in *C. jejuni*, we subtracted the SBS spectrum of *E. coli* lineage 34 from the SBS spectrum of each of the five *C. jejuni* clusters. The elevated mutations were decomposed using the signal R package signature.tools.lib v2.1.2 (*4*) into the set of bacterial DNA repair gene signatures we extracted above.

DNA polymerase III structure modelling

To compare the structures of DNA polymerase III subunits between *P. aeruginosa* and *B. cenocepacia*, we carried out structural modelling. Protein sequences were obtained for each subunit in each species from UniProt (*32*) (**table S8**). Homology models were built using SWISS-MODEL online server (*33*) for all subunits except gamma, for which a local installation of AlphaFold v2.0 (*34*) was used to build models due to sequence coverage below 95% (**table S8**). We selected the top scoring models for structural analysis in each case. ChimeraX (*35*) was used for the calculation of electrostatic surfaces, structural alignment and visualisation of predicted models. Template structures for alignment are shown in **table S8**.

De novo signature extraction

Mutational signatures were extracted from each of 14 datasets containing SBS spectra from multiple clades within a species or genus (**table S3**). We used SigProfilerExtractor v1.1.0 (*19*) which uses nonnegative matrix factorization (NMF) to split a matrix of mutation counts into underlying matrices of mutational signatures and their activities within each input SBS spectrum. The number of signatures is initially set to one and is increased up to a maximum of 25. We identified the optimal number of signatures for each dataset through comparison of the average signature stability (reflecting how well supported the signatures are within the data), mean sample cosine distance (reflecting how well the signatures fit the input SBS spectra) and individual signature stabilities.

We identified cases where the same mutational signature was extracted from multiple datasets by carrying out a hierarchical clustering of all extracted mutational signatures based on cosine distances (calculated as one minus cosine similarity). Signatures were combined if they clustered at cosine distance <0.05, corresponding to cosine similarity >0.95. Where signatures were combined, the final mutational signature was calculated as the mean of the combined signatures. The majority of combined signatures were extracted from taxonomically-nested datasets, with the exception of Bacteria_SBS15 which was extracted from several non-nested species and genus datasets within Bacillota.

The activity of each signature within each SBS spectrum was calculated as the maximum proportion of mutations assigned by SigProfilerExtractor to the signature within any extraction in which the SBS spectrum was included and the signature was extracted.

<u>Testing the impact of pathogen niche on mutational spectra</u>

We compared SBS spectra across lung-infecting and environmental clades of *Mycobacteria* and *Burkholderia* through principal component analysis (PCA) of: 1) proportions of the 96 contextual mutation in SBS spectra, 2) proportions of the six mutation types in SBS spectra and 3) proportions of the 16 contextual mutations within each mutation type. To directly compare the SBS spectra of closely related pairs of lung-infecting and environmental clades, we subtracted the SBS spectrum of the environmental clade from that of the lung-infecting clade. The mutations elevated within each clade were decomposed into potential underlying inputs using signal at https://signal.mutationalsignatures.com/ (date last accessed 24/06/2022) (*4*). We decomposed mutations elevated in environmental clades into the full set of Environmental Mutagen Signatures, excluding those associated with drug therapy which are unlikely to operate on environmental bacteria. Known and hypothesised lung-infecting clades were decomposed into the full set of lung signatures.

We carried out a targeted NMF decomposition on the full set of SBS spectra from *Mycobacteria* and *Burkholderia* using SigProfilerExtractor v1.1.0 (*19*). The presence of six signatures was identified as optimum by SigProfilerExtractor and exhibited high average signature stability (0.96 out of maximum 1) and low mean cosine distance between the input and reconstructed SBS spectra (0.016 corresponding to a mean cosine similarity of 0.984). To determine whether any of these signatures are likely niche-associated, we compared the proportion of mutations assigned to SBS spectra from the lung with SBS spectra from the environment and identified one signature that consistently exhibits a higher proportion within lung SBS spectra. We therefore named this signature Bacteria_Lung1.

We additionally examined known enteric and invasive clades from multiple *Salmonella* serovars. We split the ST313 SBS spectrum calculated in the 84 clade dataset into known enteric and invasive clades and additionally reconstructed SBS spectra for *S.* Typhimurium sequence types ST19 and ST34 (both enteric); the Global Enteric (enteric), West Africa (invasive) and East and Central Africa (invasive) clades within *S.* Enteritidis; 3 clades within *S.* Typhi (invasive); 3 clades within *S.* Paratyphi A (invasive); and 1 clade within *S.* Paratyphi C (invasive) (**table S1,** accession numbers listed in t**able S7**). To increase the number of enteric SBS spectra, and therefore increase sensitivity of downstream analyses, we ran FastBAPS v1.0.6 (*36*) on *S.* Typhimurium ST19 and the enteric clade of *S.* Typhimurium

ST313 and extracted SBS spectra of three large FastBAPS clusters in each case. Relationships between SBS spectra were inferred through hierarchical clustering of cosine distances. We compared mutation proportions in enteric and invasive clades by subtracting the SBS spectrum of the invasive clade from that of the enteric clade.

We tested whether we could predict known *Mycobacteria* and *Salmonella* niches through a set of leave-one-clade-out classifiers: k-nearest neighbours with one, two and three neighbours and a support vector machine. Each classifier was trained on cosine distances between SBS spectra, the proportion of the 96 contextual mutations and the proportion of the six mutation types, in each case excluding one random SBS spectrum. The niche of this SBS spectrum was then predicted and compared with its known niche. Only *Mycobacteria* datasets with a known niche were included in the classifiers and were labelled as lung or environmental. *Salmonella* datasets were classified into enteric and invasive niches. We assessed the ability of each classifier to identify each niche through sensitivity, specificity and area under the receiver operating characteristic curve (**fig. S16, table S5**).

We compared the DBS spectra calculated by MutTui v1.1.10 between the skin bacteria *C. acnes* and *S. epidermidis*. The DBS spectra of *S. epidermidis* phylogenetic groups A, B and C were combined for this analysis.

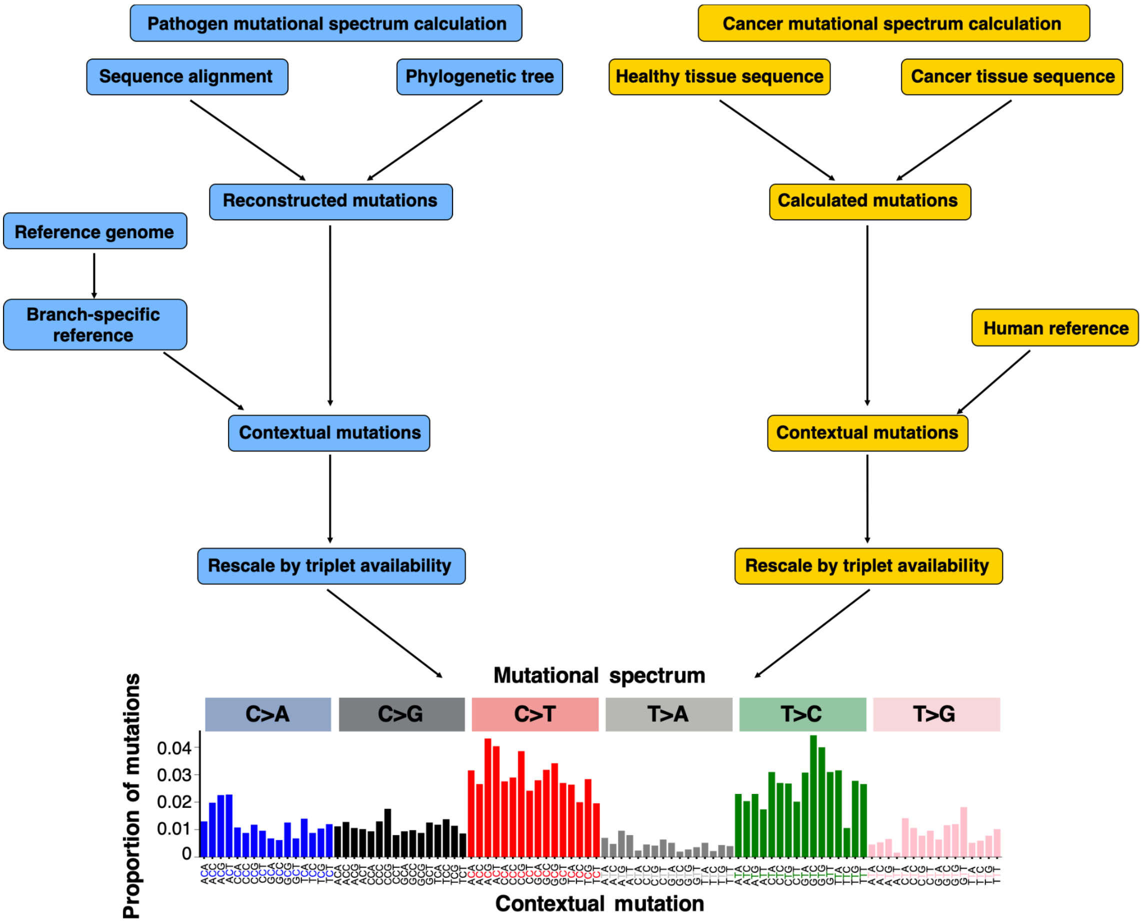


**Fig. S1. Summary of the pipeline used to calculate pathogen SBS spectra and comparison to cancer SBS spectrum calculation.** To calculate pathogen SBS spectra, we initially reconstruct directional mutations from an alignment of genetic sequences and a phylogenetic tree. The context of each mutation is identified from a reference genome that is updated at each phylogenetic node to incorporate mutations that have occurred preceding the node within the phylogenetic tree, thereby enabling identification of the context of each mutation within the branch on which it occurred. Contextual mutations are rescaled by triplet availability within the reference genome to enable comparison between bacteria. We have implemented pathogen mutational spectrum calculation in the open-source software tool MutTui (https://github.com/chrisruis/MutTui).

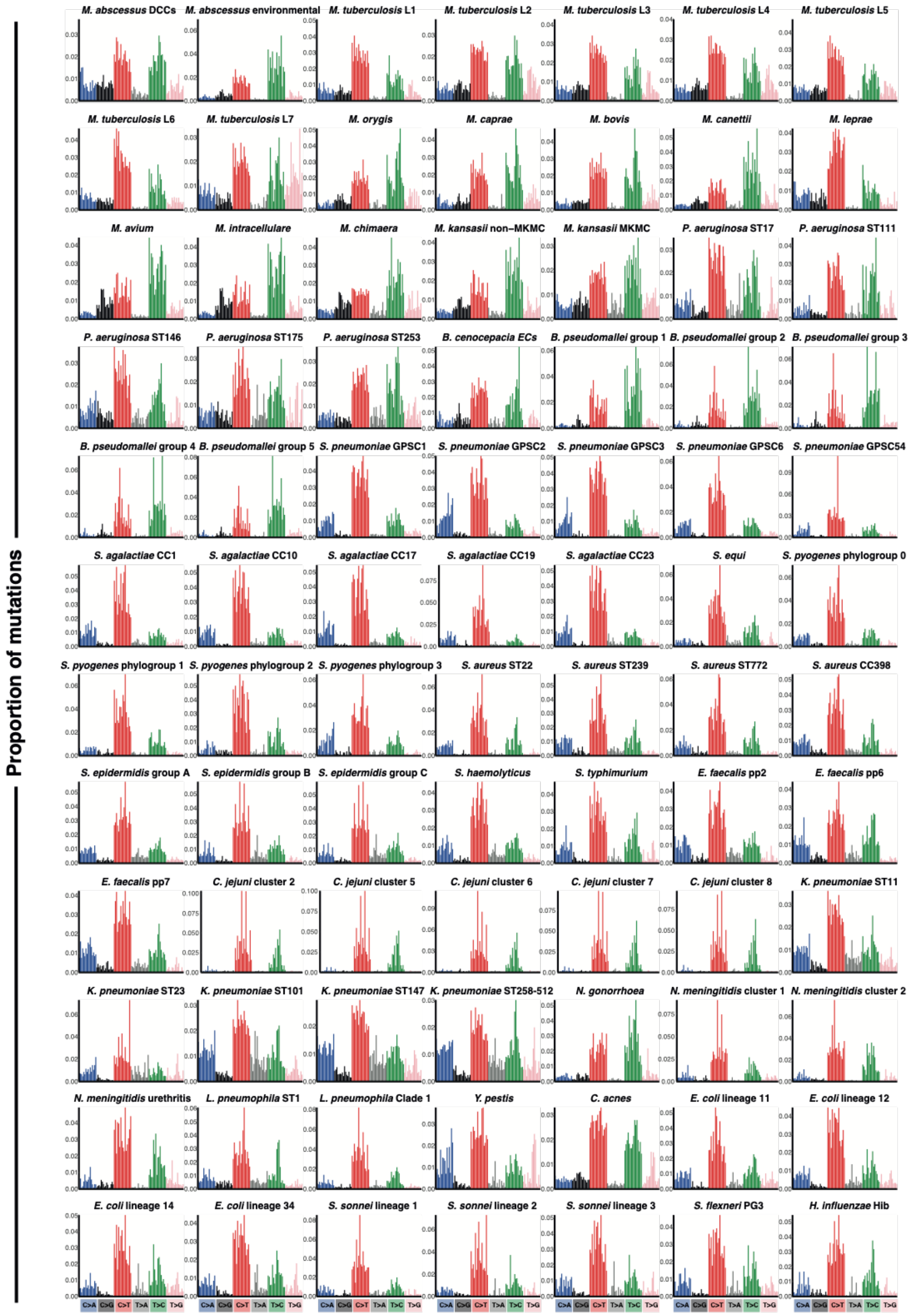


**Fig. S2. SBS spectra reconstructed from 84 phylogenetic groups across 31 bacterial species.** We assembled datasets containing whole genome sequence alignments and phylogenetic trees from previous publications and reconstructed SBS spectra as described in **fig. S1**.

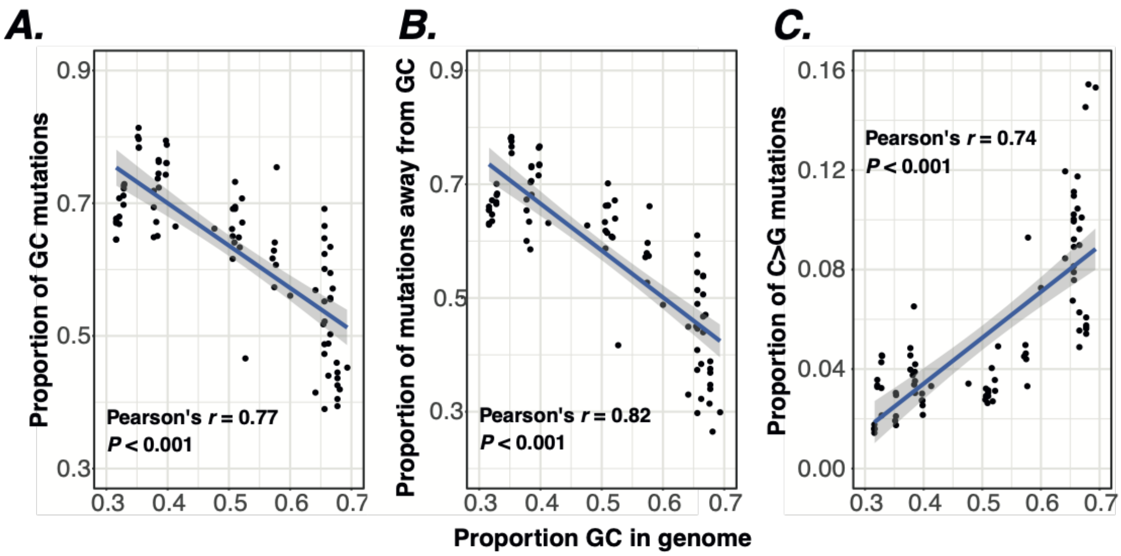


**Fig. S3. Correlation between genomic G+C content and GC base mutations.** Correlation between the proportion of G and C nucleotides in the genome with (**A**) the proportion of mutations of GC base pairs (including C>A, C>G and C>T), (**B**) the proportion of mutations from a GC base pair to an AT or TA base pair (including C>A and C>T) and (**C**) the proportion of C>G mutations. All 84 SBS spectra in **fig. S2** are included.

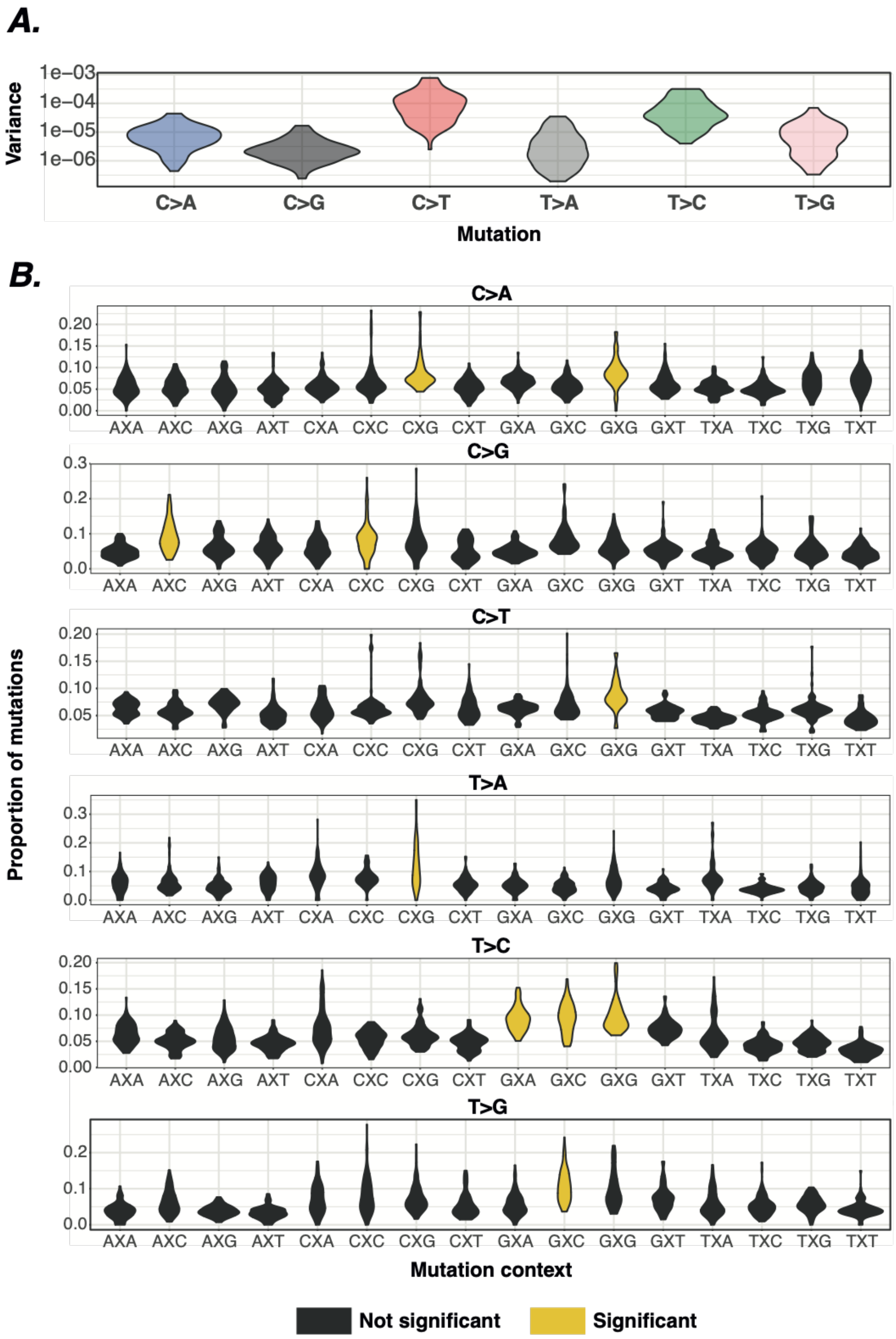


**Fig. S4. Summaries of mutation contexts across datasets.** (**A**) Distribution of variance of the 16 contextual mutation proportions within each mutation type across the 84 SBS spectra. C>T and T>C exhibit elevated context-specificity (Tukey HSD corrected two-way ANOVA *P* < 0.001). (**B**) Distribution of contextual mutation proportions within each mutation type across the 84 SBS spectra. Significant contexts exhibit median context proportion at least 2.5 median average deviations away from the median proportion across all contexts within the mutation type.

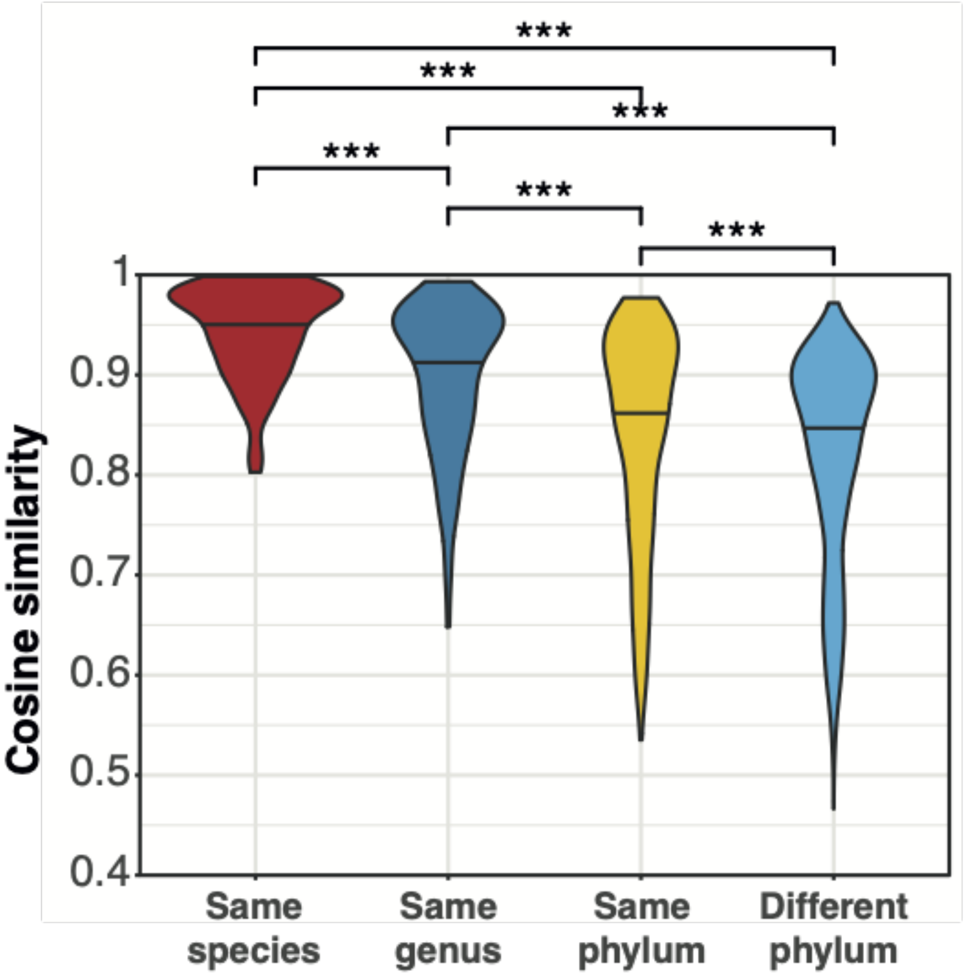


**Fig. S5. Correlation between phylogenetic relatedness and SBS spectrum similarity.** We calculated the cosine similarity between all SBS spectrum pairs and split based on taxonomic relationship. Cosine similarities are only placed in the lowest taxonomic match, for example if spectra are from the same species, they will not be included in the same genus comparisons. Significance was calculated using Tukey HSD corrected two-way ANOVA between the distributions, *** - $P < 0.001$.

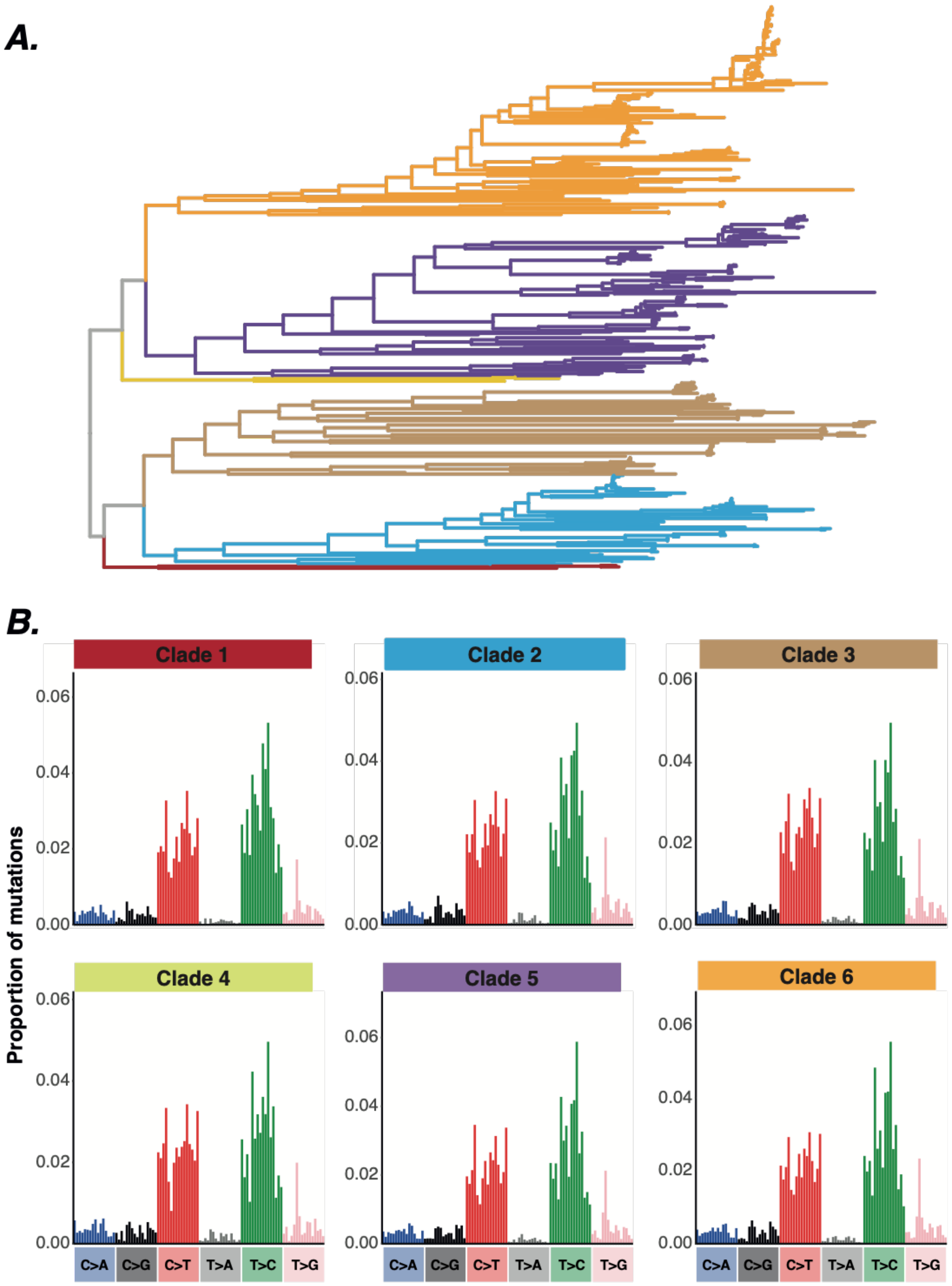


**Fig. S6. Comparison of SBS spectra between *Neisseria gonorrhoeae* clades.** (**A**) Phylogenetic tree of 412 *N. gonorrhoeae* isolates used to reconstruct the SBS spectrum. The tree is coloured by clade based on phylogenetic clustering. The *N. gonorrhoeae* SBS spectrum included in other analyses was calculated across the complete tree. (**B**) SBS spectrum of each of the six clades highlighted in panel **A**. SBS spectra are highly similar between clades, cosine similarity >0.98 between all pairs.

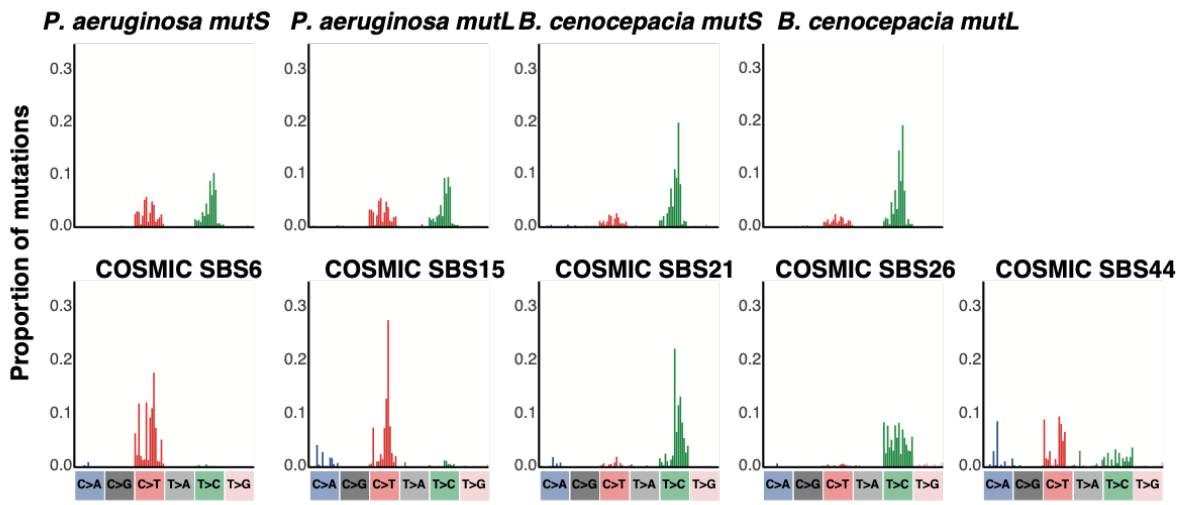


**Fig. S7. Comparison of MMR signatures between bacteria and humans.** The top row shows the MMR signatures extracted from bacterial hypermutator lineages, as in **Fig. 2B**. The bottom row shows the five COSMIC signatures currently associated specifically with defective MMR in humans (*1*, *8*, *12*).

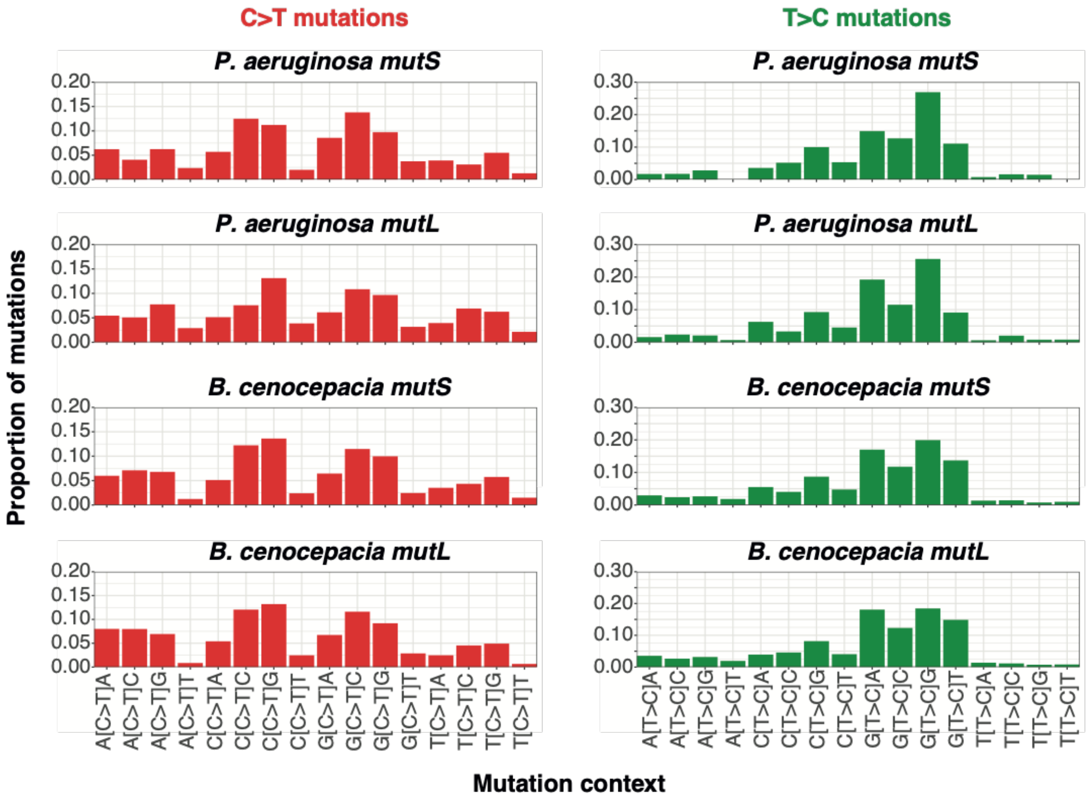


**Fig. S8. Comparison of contextual C>T and T>C mutations between mismatch repair hypermutators in *P. aeruginosa* and *B. cenocepacia*.** The proportion of each contextual mutation within C>T or T>C is shown. Cosine similarity between contextual mutation proportions within C>T or within T>C >0.95 between all pairs.

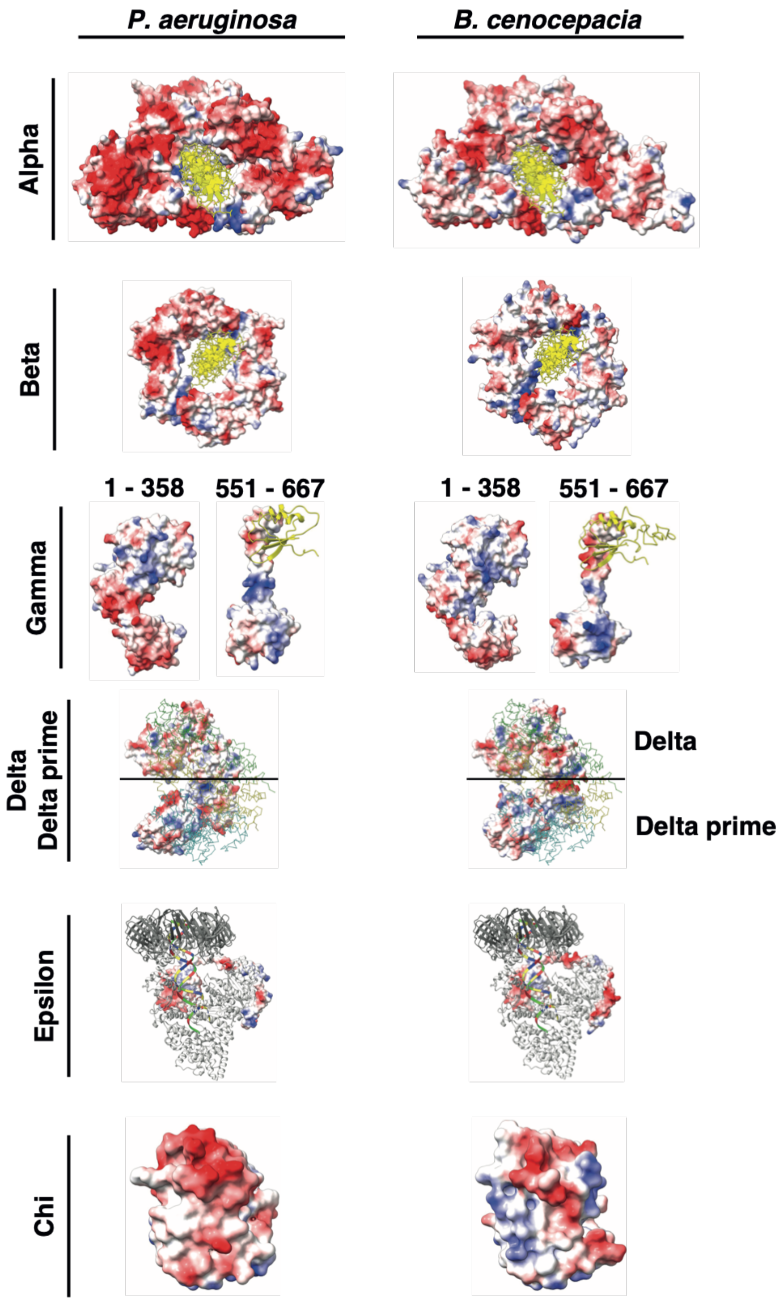


**Fig. S9. Structural modelling of DNA polymerase III.** Predicted models of each subunit of DNA polymerase III in *P. aeruginosa* and *B. cenocepacia* are shown as electrostatic surfaces with negative charges shown in red and positive charges in blue. Alpha and beta subunit predicted models are superposed on the crystal structure of *E. coli* replicative DNA polymerase complex (PDB ID 5FKV) that contains bound DNA (shown in yellow as stick representation). Residues with predicted local distance difference test (pLDDT) score greater than 70 are shown for the gamma subunit. Numbers above the structures indicate included residues. Gamma subunit models include the c-terminal residues of the alpha subunit (yellow ribbon) that interact with the gamma subunit, inferred by superposing the respective subunits onto the cryo EM structure of the *E. coli* replicative DNA polymerase complex (PDB ID 5FKV). Horizontal lines in the delta and delta prime models separate the interface between delta (above the line) and delta prime (below the line) subunits; the interaction interface is inferred by superposing the homology models onto the respective subunits of the crystal structure of the *E.coli* clamp loader complex (PDB ID 1XXI; gamma subunit chains B, C and D shown as c-alpha trace in green, yellow and cyan respectively). Predicted epsilon subunit models are superposed onto the cryo EM structure of the *E. coli* replicative DNA polymerase complex (PDB ID 5FKW; alpha and beta subunits shown as ribbon in light grey and dark grey, respectively, bound DNA shown as coloured ribbon). Overall DNA polymerase III subunit structures are similar between *P. aeruginosa* and *B. cenocepacia* but electrostatic surfaces are distinct.

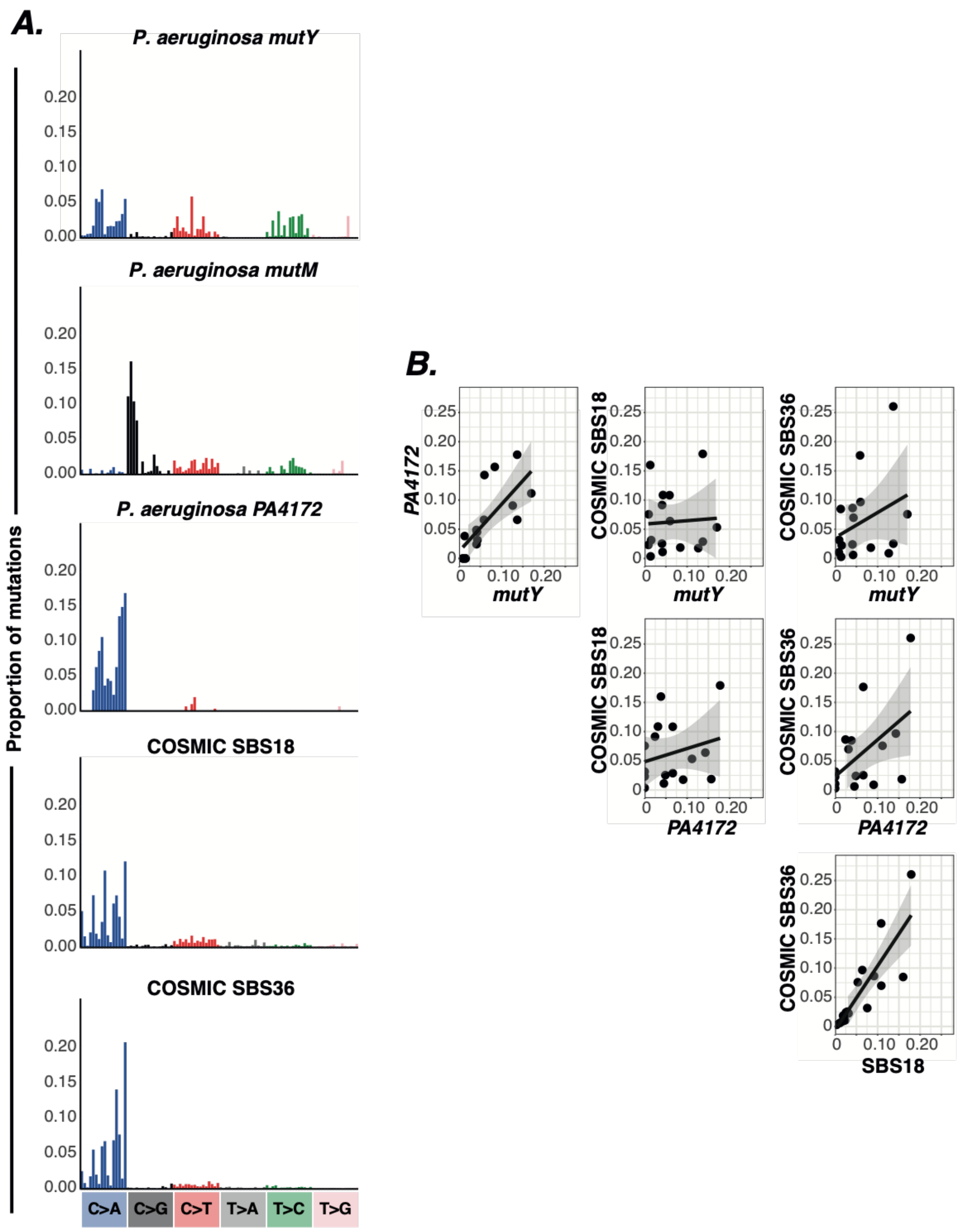


**Fig. S10. Comparison of GO signatures between bacteria and humans. (A)** Mutational signatures of GO genes extracted from bacteria and COSMIC signatures associated with GO genes in humans (*9*). (**B**) Regression of the proportion of the 16 contextual mutations within C>A between gene signatures that exhibit C>A mutations. The correlation is significant between *P. aeruginosa mutY* and *P. aeruginosa PA4172* (Pearson's $r = 0.73$; 95% CI: 0.37, 0.9; Benjamini-Hochberg corrected $P = 0.005$) and between COSMIC SBS18 and COSMIC SBS36 (Pearson's $r = 0.84$; 95% CI: 0.59, 0.94; Benjamini-Hochberg corrected $P < 0.001$).

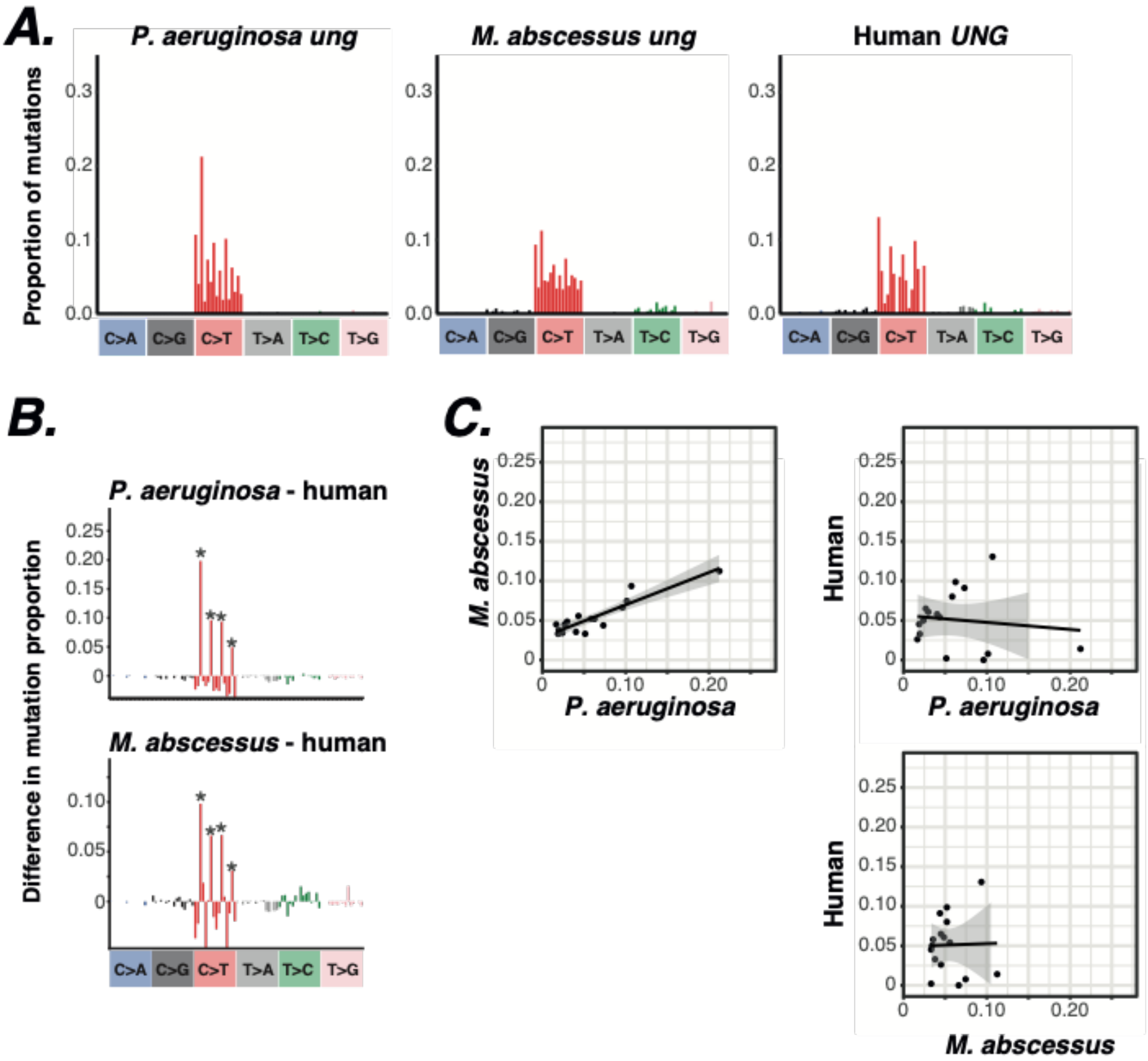


**Fig. S11. Comparison of *ung* signatures between bacteria and humans. (A)** Mutational signatures of *ung* gene hypermutator lineages extracted from bacteria and *in vitro UNG* knockout in human cells (*9*). (**B**) Subtraction of the human *UNG* knockout signature from the bacterial *ung* signatures. Asterisks indicate CpG contexts. (**C**) Regression of the proportion of the 16 contextual mutations within C>T between gene signatures. The correlation is significant between *P. aeruginosa ung* and *M. abscessus ung* (Pearson's $r$ = 0.91; 95% CI: 0.75, 0.97; Benjamini-Hochberg corrected $P < 0.001$).

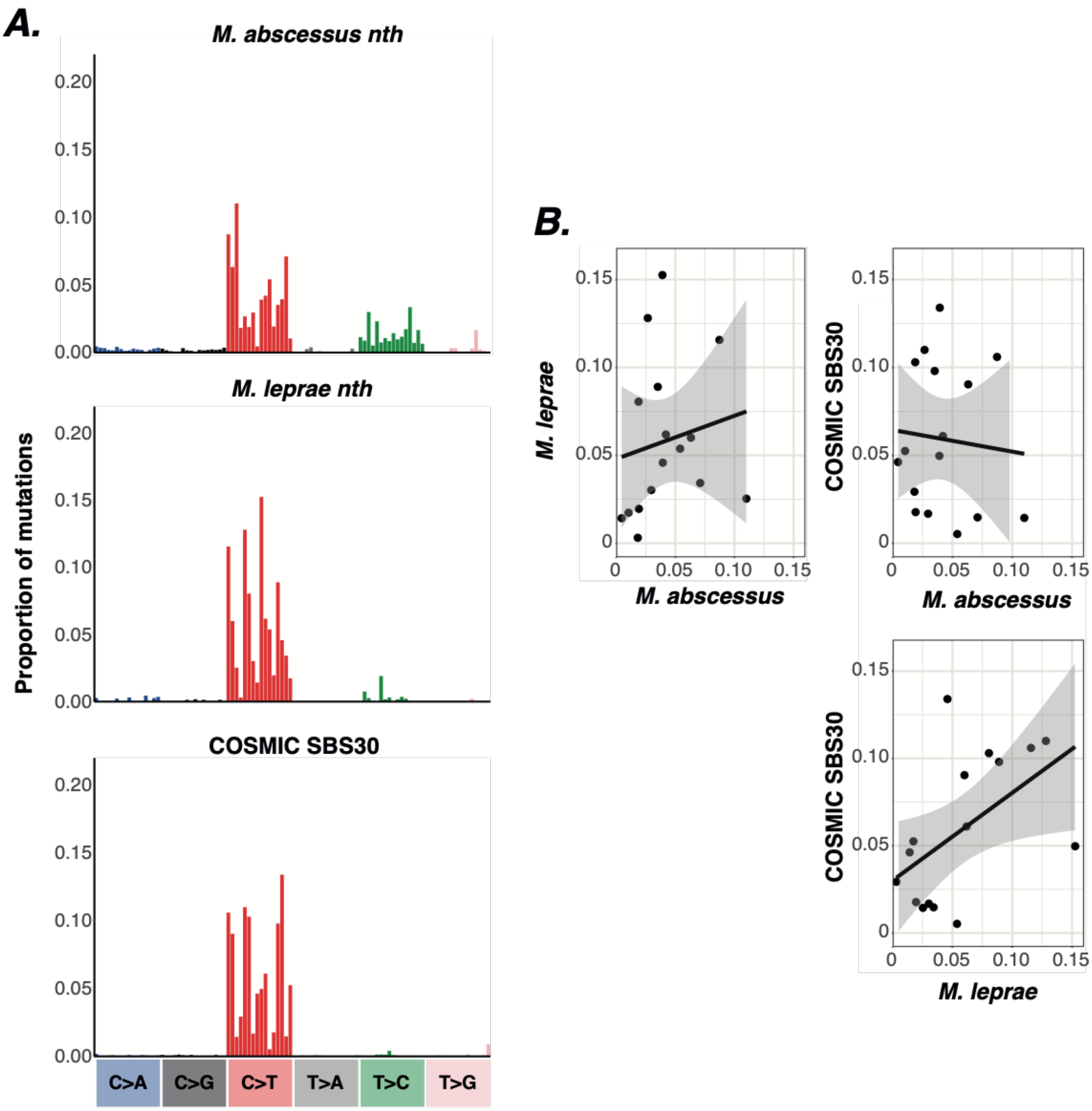


**Fig. S12. Comparison of *nth* signatures between bacteria and humans. (A)** Mutational signatures of *nth* gene hypermutator lineages extracted from bacteria and COSMIC SBS30 which is associated with *in vitro* knockout of the *nth* homologue *NTHL1* in human cells (*9*). (**B**) Regression of the proportions of the 16 contextual mutations within C>T between gene signatures. None of the correlations are statistically significant.

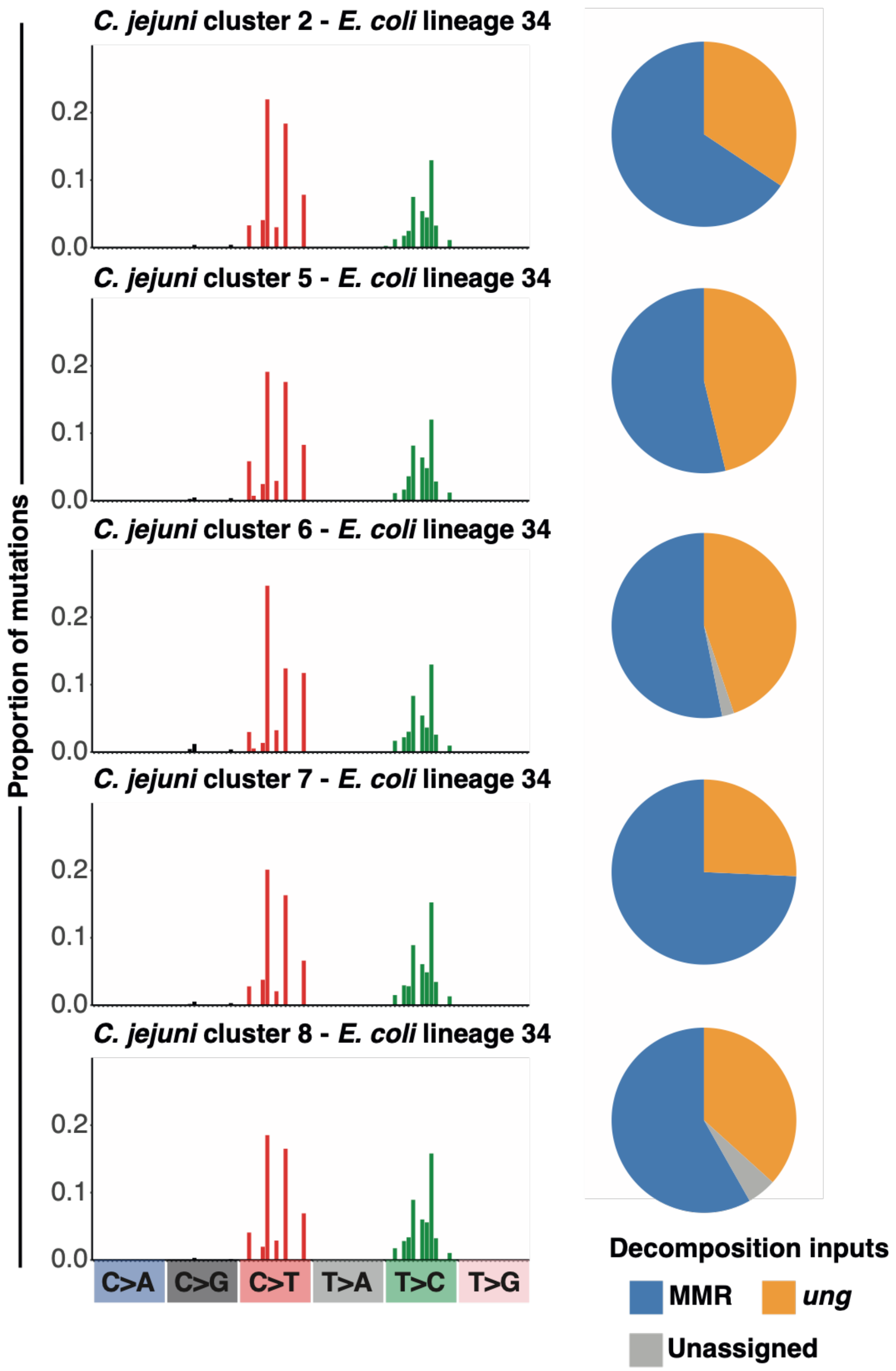


**Fig. S13. Signatures of DNA repair deficiency in *C. jejuni*.** Spectrum plots show the SBS mutations elevated in the respective *C. jejuni* cluster compared with *E. coli* lineage 34 which occupies a similar niche. Due to the shared niche, these elevated mutations are likely the result of processes ongoing within *C. jejuni*. Pie charts show the proportion of elevated mutations assigned to the respective bacterial DNA repair signature in a decomposition analysis into the full set of extracted bacterial DNA repair signatures in **Fig. 2B-D**.

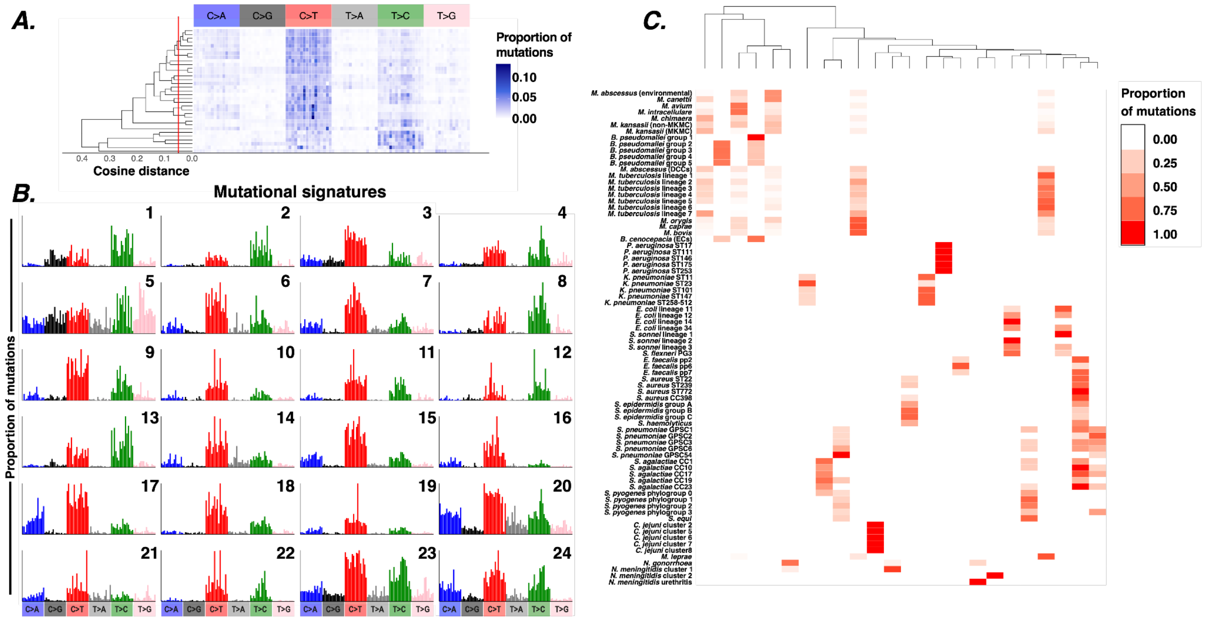


**Fig. S14. Mutational signatures extracted from decomposition analysis of datasets of species and genus SBS spectra. (A)** Hierarchical clustering of the 33 mutational signatures extracted from 13 extraction datasets containing SBS spectra from clades within a species or within a genus. Signatures were combined if they cluster at cosine similarity of 0.95, shown by the red vertical line. Based on this, the 33 extracted mutational signatures were collapsed to 24 final mutational signatures. The heatmap shows the proportion of contextual mutations within each mutational signature. (**B**) SBS spectra for the 24 extracted mutational signatures showing the proportion of each contextual mutation. (**C**) Activity of each extracted signature within each SBS spectrum included in decomposition analysis, calculated as the maximum proportion of mutations assigned to the signature across decompositions. The dendrogram shows hierarchical clustering of the 24 composite signatures.

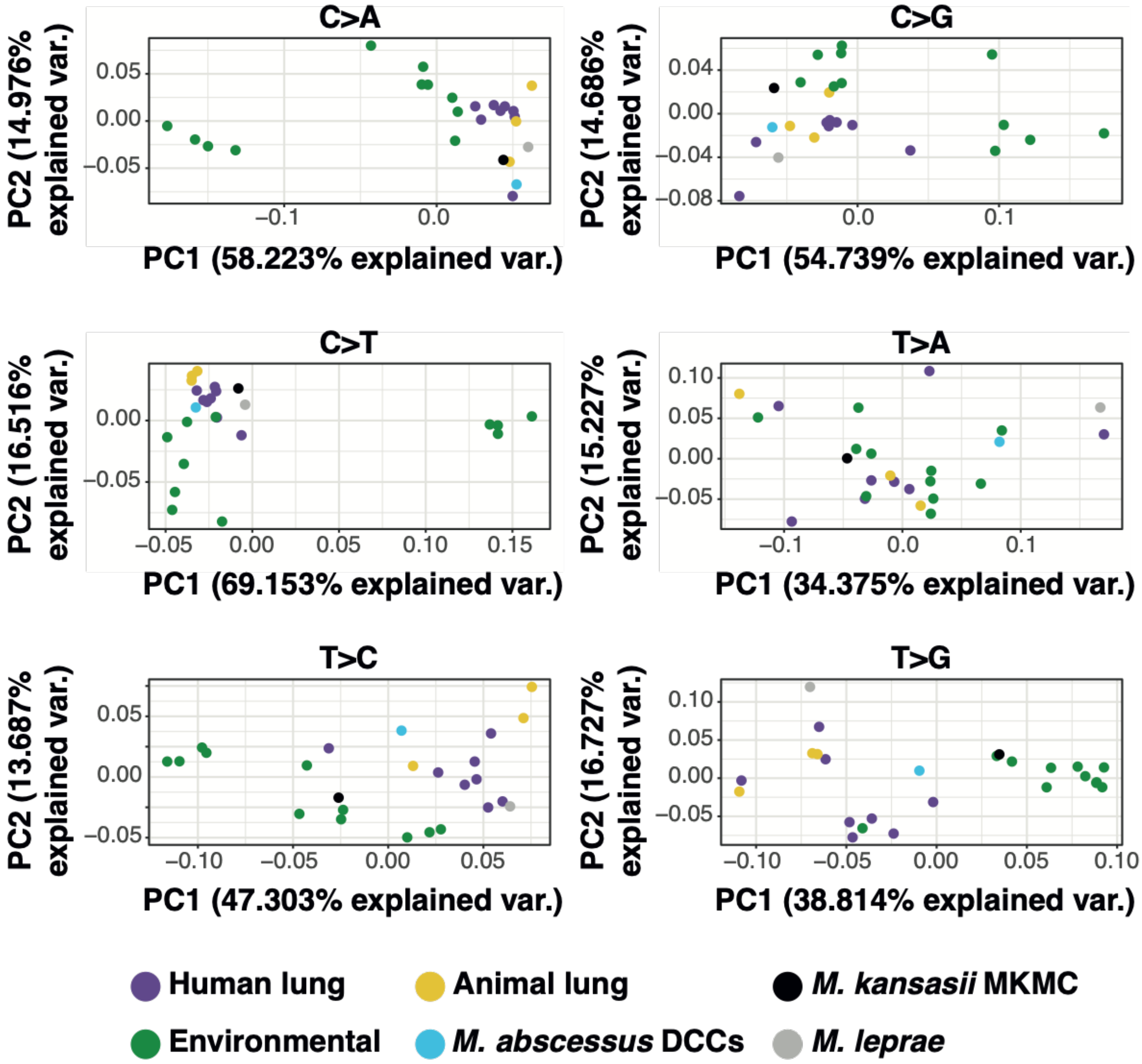


**Fig. S15. Contextual mutation comparisons between *Mycobacteria* and *Burkholderia*.** PCA of contextual mutation proportions within each mutation type across *Mycobacteria* and *Burkholderia* SBS spectra.

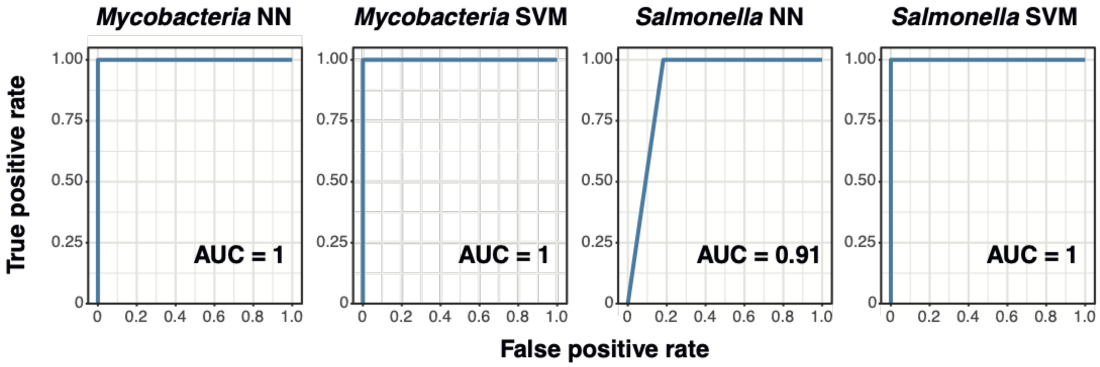


**Fig. S16. ROC curves for niche predictors within *Mycobacteria* and *Salmonella*.** We developed a set of leave-one-out classifiers to predict niche based on SBS spectrum. Each included spectrum has a known replication niche, either environment or lung for included *Mycobacteria* and either enteric or invasive for included *Salmonella*. The shown classifiers predict niche based on cosine similarity between the left out spectrum and remaining spectra using either three nearest neighbours (NN) or a support vector machine (SVM). AUC: area under curve.

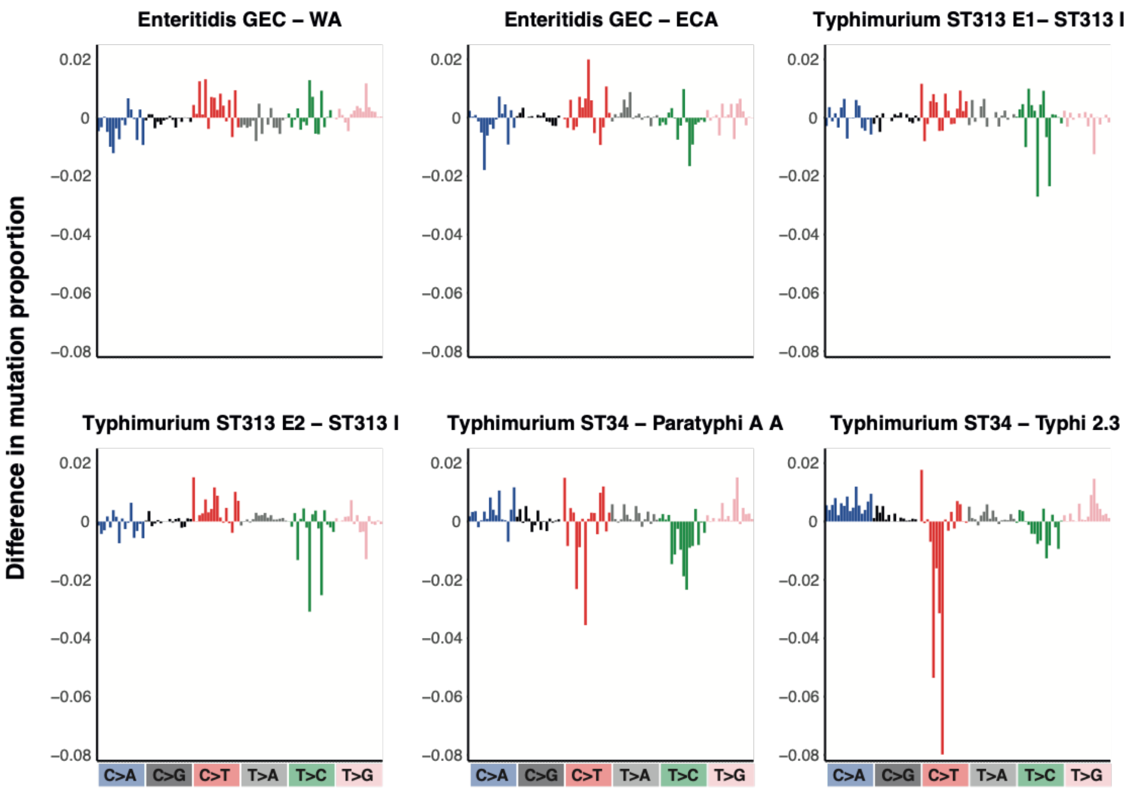


**Fig. S17. Comparison of enteric and invasive *Salmonella* SBS spectra.** The proportion of each contextual mutation in the respective invasive spectrum was subtracted from that in the respective enteric spectrum. Mutations above zero are more common in the enteric spectrum, mutations below zero are more common in the invasive spectrum. GEC = Global Enteric clade, WA = West Africa clade, ECA = East and Central Africa clade.

**Table S1. (separate file)** Datasets used for reconstruction of SBS and DBS mutational spectra.

**Table S2. (separate file)** SBS spectra calculated from the 84 bacterial clades.

**Table S3. (separate file)** Datasets used for NMF signature extraction.

**Table S4. (separate file)** Bacteria SBS signatures and the datasets they were extracted from.

**Table S5. (separate file)** Summary of niche classifiers.

**Table S6. (separate file)** SBS spectra calculated from enteric and invasive *Salmonella* lineages.

**Table S7. (separate file)** Accession numbers of samples used in mutational spectrum datasets.

**Table S8. (separate file)** Modelling of DNA polymerase III subunits in *Pseudomonas aeruginosa* and *Burkholderia cenocepacia*.
